# Rare coding variants in 35 genes associate with circulating lipid levels – a multi-ancestry analysis of 170,000 exomes

**DOI:** 10.1101/2020.12.22.423783

**Authors:** George Hindy, Peter Dornbos, Mark D. Chaffin, Dajiang J. Liu, Minxian X. Wang, Margaret Sunitha Selvaraj, David Zhang, Joseph Park, Carlos A. Aguilar-Salinas, Lucinda Antonacci-Fulton, Diego Ardissino, Donna K. Arnett, Stella Aslibekyan, Gil Atzmon, Christie M. Ballantyne, Francisco Barajas-Olmos, Nir Barzilai, Lewis C. Becker, Lawrence F. Bielak, Joshua C. Bis, John Blangero, Eric Boerwinkle, Lori L. Bonnycastle, Erwin Bottinger, Donald W. Bowden, Matthew J. Bown, Jennifer A. Brody, Jai G. Broome, Noël P. Burtt, Brian E. Cade, Federico Centeno-Cruz, Edmund Chan, Yi-Cheng Chang, Yii-Der I. Chen, Ching-Yu Cheng, Won Jung Choi, Rajiv Chowdhury, Cecilia Contreras-Cubas, Emilio J. Córdova, Adolfo Correa, L Adrienne Cupples, Joanne E. Curran, John Danesh, Paul S. de Vries, Ralph A. DeFronzo, Harsha Doddapaneni, Ravindranath Duggirala, Susan K. Dutcher, Patrick T. Ellinor, Leslie S. Emery, Jose C. Florez, Myriam Fornage, Barry I. Freedman, Valentin Fuster, Ma. Eugenia Garay-Sevilla, Humberto García-Ortiz, Soren Germer, Richard A. Gibbs, Christian Gieger, Benjamin Glaser, Clicerio Gonzalez, Maria Elena Gonzalez-Villalpando, Mariaelisa Graff, Sarah E Graham, Niels Grarup, Leif C. Groop, Xiuqing Guo, Namrata Gupta, Sohee Han, Craig L. Hanis, Torben Hansen, Jiang He, Nancy L. Heard-Costa, Yi-Jen Hung, Mi Yeong Hwang, Marguerite R. Irvin, Sergio Islas-Andrade, Gail P. Jarvik, Hyun Min Kang, Sharon L.R. Kardia, Tanika Kelly, Eimear E. Kenny, Alyna T. Khan, Bong-Jo Kim, Ryan W. Kim, Young Jin Kim, Heikki A. Koistinen, Charles Kooperberg, Johanna Kuusisto, Soo Heon Kwak, Markku Laakso, Leslie A. Lange, Jiwon Lee, Juyoung Lee, Seonwook Lee, Donna M. Lehman, Rozenn N. Lemaitre, Allan Linneberg, Jianjun Liu, Ruth J.F. Loos, Steven A. Lubitz, Valeriya Lyssenko, Ronald C.W. Ma, Lisa Warsinger Martin, Angélica Martínez-Hernández, Rasika A. Mathias, Stephen T. McGarvey, Ruth McPherson, James B. Meigs, Thomas Meitinger, Olle Melander, Elvia Mendoza-Caamal, Ginger A. Metcalf, Xuenan Mi, Karen L. Mohlke, May E. Montasser, Jee-Young Moon, Hortensia Moreno-Macías, Alanna C. Morrison, Donna M. Muzny, Sarah C. Nelson, Peter M. Nilsson, Jeffrey R. O’Connell, Marju Orho-Melander, Lorena Orozco, Colin N.A. Palmer, Nicholette D. Palmer, Cheol Joo Park, Kyong Soo Park, Oluf Pedersen, Juan M. Peralta, Patricia A. Peyser, Wendy S. Post, Michael Preuss, Bruce M. Psaty, Qibin Qi, DC Rao, Susan Redline, Alexander P. Reiner, Cristina Revilla-Monsalve, Stephen S. Rich, Nilesh Samani, Heribert Schunkert, Claudia Schurmann, Daekwan Seo, Jeong-Sun Seo, Xueling Sim, Rob Sladek, Kerrin S. Small, Wing Yee So, Adrienne M. Stilp, E Shyong Tai, Claudia H.T. Tam, Kent D. Taylor, Yik Ying Teo, Farook Thameem, Brian Tomlinson, Michael Y. Tsai, Tiinamaija Tuomi, Jaakko Tuomilehto, Teresa Tusié-Luna, Rob M. van Dam, Ramachandran S. Vasan, Karine A. Viaud Martinez, Fei Fei Wang, Xuzhi Wang, Hugh Watkins, Daniel E. Weeks, James G. Wilson, Daniel R. Witte, Tien-Yin Wong, Lisa R. Yanek, AMP-T2D-GENES, Myocardial Infarction Genetics Consortium, NHLBI Trans-Omics for Precision Medicine (TOPMed) Consortium, NHLBI TOPMed Lipids Working Group, Sekar Kathiresan, Daniel J. Rader, Jerome I. Rotter, Michael Boehnke, Mark I. McCarthy, Cristen J. Willer, Pradeep Natarajan, Jason A. Flannick, Amit V. Khera, Gina M. Peloso

## Abstract

Large-scale gene sequencing studies for complex traits have the potential to identify causal genes with therapeutic implications. We performed gene-based association testing of blood lipid levels with rare (minor allele frequency<1%) predicted damaging coding variation using sequence data from >170,000 individuals from multiple ancestries: 97,493 European, 30,025 South Asian, 16,507 African, 16,440 Hispanic/Latino, 10,420 East Asian, and 1,182 Samoan. We identified 35 genes associated with circulating lipid levels. Ten of these: *ALB*, *SRSF2*, *JAK2, CREB3L3*, *TMEM136*, *VARS*, *NR1H3*, *PLA2G12A*, *PPARG* and *STAB1* have not been implicated for lipid levels using rare coding variation in population-based samples. We prioritize 32 genes identified in array-based genome-wide association study (GWAS) loci based on gene-based associations, of which three: *EVI5, SH2B3*, and *PLIN1*, had no prior evidence of rare coding variant associations. Most of the associated genes showed evidence of association in multiple ancestries. Also, we observed an enrichment of gene-based associations for low-density lipoprotein cholesterol drug target genes, and for genes closest to GWAS index single nucleotide polymorphisms (SNP). Our results demonstrate that gene-based associations can be beneficial for drug target development and provide evidence that the gene closest to the array-based GWAS index SNP is often the functional gene for blood lipid levels.

## Introduction

Blood lipid levels are heritable complex risk factors for atherosclerotic cardiovascular diseases.^1^ Array-based genome-wide association studies (GWAS) have identified >400 loci as associated with blood lipid levels, explaining 9-12% of the phenotypic variance of lipid traits.^2–8^ These studies have identified mostly common (minor allele frequency (MAF)>1%) noncoding variants with modest effect and helped define the causal roles of different lipid fractions in cardiovascular disease.^9–13^ Despite these advances, the mechanisms and causal genes for most of the identified variants and loci have not yet been determined.

Conventional GWAS with array-derived or imputed common variants are unlikely to directly implicate causal genes, while genetic association studies testing rare variants in coding regions have this potential. Advances in next generation sequencing over the last decade have facilitated increasingly larger studies with improved power to detect associations of rare variants with complex diseases and traits.^14; 15^ However, most exome sequencing studies to date have been insufficiently powered for rare variant discovery; for example, Flannick et al. estimated that it would require 75,000 to 185,000 sequenced cases of type 2 diabetes (T2D) to detect associations at known drug target genes at exome-wide significance.^15^

Identifying rare variants with impact on protein function has helped elucidate biological pathways underlying dyslipidemia and atherosclerotic diseases such as coronary artery disease (CAD).^14; 16–25^ Successes using this approach have led to the development of novel therapeutic targets to modify blood lipid levels and lower risk of atherosclerotic diseases.^26; 27^ The vast majority of participants in these studies have been of European ancestry, highlighting the need for more diverse study sample. Such diversity can identify associated variants absent or present at very low frequencies in European populations and help implicate new genes with generalizability extending to all populations.

We have assembled exome sequence data from >170,000 individuals across multiple ancestries and systematically tested the association of rare variants in each gene with six circulating lipid phenotypes: low-density lipoprotein cholesterol (LDL-C), high density lipoprotein cholesterol (HDL-C), non-HDL-C, total cholesterol (TC), triglycerides (TG), and the ratio of TG to HDL-C (TG:HDL). We find 35 genes associated with blood lipid levels, show evidence of gene-based signals in array-based GWAS loci, show enrichment of lipid gene-based associations in LDL-C drug targets and genes in close proximity with GWAS index variants, and test lipid genes for association with CAD, T2D, and liver enzymes.

## Subjects and Methods

### Study Overview

Our study samples were derived from four major data sources with exome or genome sequence data and blood lipid levels: CAD case-control studies from the Myocardial Infarction Genetics Consortium (MIGen, n = 44,208) and a UKB nested case-control study of CAD (n = 10,689); T2D cases-control studies from the AMP-T2D-GENES exomes (n = 32,486); population-based studies from the TOPMed project freeze 6a data (n = 44,101) restricted to the exome, and the UKB first tranche of exome sequence data (n = 40,586) (see **Supplementary Note**). Informed consent was obtained from all subjects and committees approving the studies are available in the supplement.

Within each data source, individuals were excluded if they failed study-specific sequencing quality metrics, lacked lipid phenotype data, or were duplicated in other sources. We additionally removed first- and second-degree relatives across data sources while we kept relatives within each data source since we were able to adjust for relatedness within each data source using kinship matrices in linear mixed models. If samples from the same study were present in different data sources, we used the samples in the data source which has the largest sample size from the study and removed the overlapping set from the other data source. For instance, samples from the Atherosclerosis Risk in Communities (ARIC) Study were removed from TOPMed and kept in MIGen which had more sequenced samples from ARIC. Similarly, samples from the Jackson Heart Study were kept in TOPMed and removed from MIGen. To obtain duplicate and kinship information across data sources we used 14,834 common (MAF>1%) and no more than weakly dependent (r^2^ < 0.2) variants using the make-king flag in PLINK v2.0. Single-variant association analyses were performed within each data source, case-status, and ancestry combination. The data were sequenced and variant calling performed separately by data source and this allowed us to look for effects by case-status and genetically-inferred and/or self-reported ancestry groups. We performed gene-based meta-analyses by combining single-variant summary statistics and covariance matrices generated from RVTESTS.^28^ We performed ancestry-specific gene-based meta-analyses by combining single-variant summary data from five major ancestries with >10,000 across all data sources: European, South Asian, African, Hispanic, and East Asian ancestries.

### Phenotypes

We studied six lipid phenotypes; total, LDL-C, HDL-C, non-HDL-C, TG and TG:HDL. TC was adjusted by dividing the value by 0.8 in individuals reporting lipid lowering medication use after 1994 or statin use at any time point. If LDL-C levels were not directly measured, then they were calculated using Friedewald equation for individuals with TG levels < 400 mg/dl using adjusted TC levels. If LDL-C levels were directly measured then, their values were divided by 0.7 in individuals reporting lipid lowering medication use after 1994 or statin use at any time point.^5^ TG and TG:HDL levels were natural logarithm transformed. Non-HDL-C was obtained by subtracting HDL-C from adjusted TC levels. Residuals for each trait in each cohort, ancestry, and case status grouping were created after adjustment for age, age^2^, sex, principal components, sequencing platform, and fasting status (when available) in a linear regression model. Residuals were then inverse-normal transformed and multiplied by the standard deviation of the trait to scale the effect sizes to the interpretable units.

### Sequencing and Quality Control

#### Myocardial Infarction Genetics Consortium (MIGen)

A set of common variants was extracted for sample quality control including relative inference, principal component analysis, and estimation of heterozygosity. SNPs on autosomes and not in low complexity regions or segmental duplications were extracted. SNPs with quality of depth (QD)> 2, call rate >98%, self-reported-race-specific Hardy-Weinberg equilibrium (HWE) p-value >1×10^-8^, Variant Quality Score Recalibration (VQSR) of PASS and MAF>1% were retained. Sample relatedness was estimated with KING and duplicate samples removed. Genetically inferred ancestry was assigned to each individual by calculating principal components jointly with 1000 Genomes phase 3 version 5 and building a 5-Nearest Neighbor classifier using the top 6 principal components. Heterozygosity was estimated within each genetic ancestry group and samples with F statistic above 0.3 were removed. Genetic sex was inferred based on high quality X-chromosome variation including variants with call rate >0.95, MAF>2%, a PASS VQSR, QD>3 if the variant is an insertion or deletion and QD>2 if it is SNP. Samples with discordant phenotypic sex and genetic sex were removed. Finally, sample quality control metrics were calculated using Hail and samples with call rate<0.9a mean depth (DP)<30 and mean genotype quality (GQ)<0.8 were excluded. A total of 44,240 samples with lipid data measurements were included after further excluding duplicates and relatives with other data sources (**Table S1**).

**Table 1.**
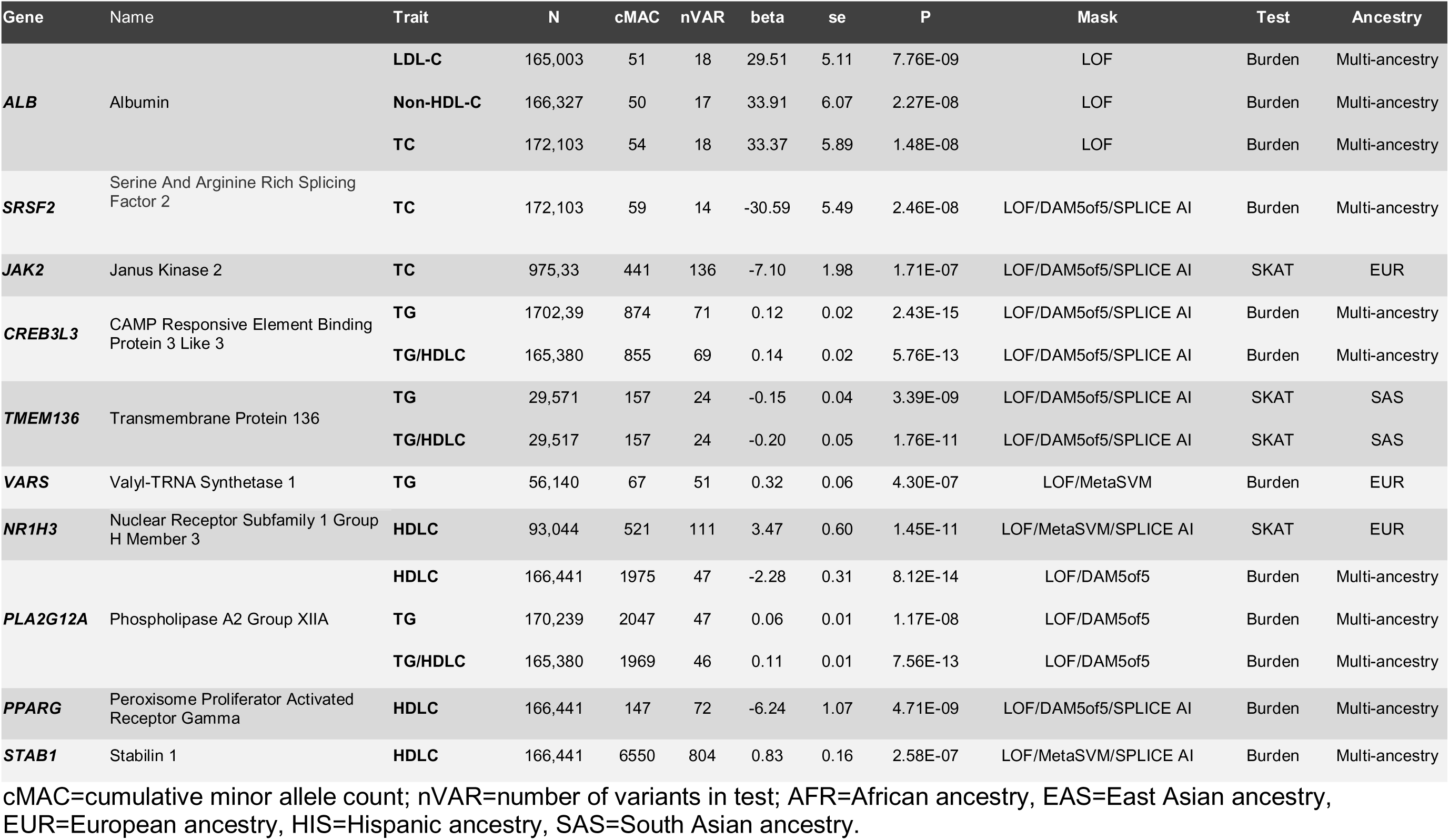
Novel Genes Associated with Blood Lipids

Variant quality control was performed amongst remaining samples and a total of 8,716,575 autosomal variants were included after removing those that fail HWE as calculated by genetic ancestry group (p-value<1×10^-8^), lie in low complexity regions or segmental duplications, with inbreeding coefficient< −0.3, are insertions or deletions with QD ≤ 3 or SNPs with QD ≤ 2 or variants where VQSR does not PASS with the exception of singletons where variants with VQSRTrancheSNP99.60to99.80 were retained.

#### Trans-Omics for Precision Medicine (TOPMed)

Whole genome sequencing at 30X mean depth was performed at one of six sequencing centers: Broad Institute of MIT and Harvard, Northwest Genomics Center, New York Genome Center, Illumina Laboratory Services, Psomagen, Inc. (formerly Macrogen USA), Baylor College of Medicine Human Genome Sequencing Center. For most studies, all individuals in the study were sequenced at the same center. Sequence reads were aligned to human genome build GRCh37 or GRCh38 at each center using similar, but not identical, processing pipelines. The resulting sequence data files were transferred from all centers to the TOPMed Informatics Research Center (IRC), where they were re-aligned to build GRCh38, using a common pipeline to produce a set of ‘harmonized’ .cram files. Processing was coordinated and managed by the ‘GotCloud’ processing pipeline. The IRC performed joint genotype calling on all samples. Quality control was performed at each stage of the process by the Sequencing Centers, the IRC, and the TOPMed Data Coordinating Center (DCC). Only samples that passed QC were included in the call set.

The two sequence quality criteria that were used to pass sequence data on for joint variant discovery and genotyping are: estimated DNA sample contamination below 3%, and fraction of the genome covered at least 10x 95% or above. DNA sample contamination was estimated from the sequencing center read mapping using software verifyBamId.^29^

The genotype used for analysis are from “freeze 6a” of the variant calling pipeline performed by the TOPMed Informatics Research Center (Center for Statistical Genetics, University of Michigan, Hyun Min Kang, Tom Blackwell and Gonçalo Abecasis). Variant detection (SNPs and indels) from each sequenced (and aligned) genome was performed by the vt discover2 software tool. The variant calling software tools are under active development; updated versions can be accessed at http://github.com/atks/vt, http://github.com/hyunminkang/apigenome, and https://github.com/statgen/topmed_variant_calling.

One individual from duplicate pairs identified by the DCC was removed, retaining the individual with lipid levels available when one did not have lipid levels. If both individuals had lipid levels, one individual was randomly selected. Individuals were excluded when their genotype determined sex did not match phenotype reported sex (n=6) and individuals <18 years old were excluded (n=865). Ancestry was defined as self-reported ancestry.

#### AMP-T2D-GENES

Sequencing and quality control were performed as previously described.^15^ Following sequencing and variant calling, we measured samples and variants according to several sequence quality metrics and excluded those that were outliers relative to the global distribution. These exclusions produced an “analysis” dataset of 45,231 individuals and 6.33M variants. We then estimated, within each ancestry, pairwise IBD values, genetic relatedness matrices (GRMs), and PCs for use in downstream association analysis. We used the IBD values to generate lists of unrelated individuals within each ancestry, excluding 2,157 individuals from an “unrelated analysis” set of 43,090 individuals (19,828 cases and 23,262 controls) and 6.29M non-monomorphic variants.

#### UK Biobank

We used two UKB datasets with exome sequence data. The first is a CAD case control study with 12,938 individuals. 29 samples were removed as they had discordant genotypes with genotyping array data, 17 showed mismatch between the reported and genetically inferred sex, 4 had excess heterozygosity and 6 had a call rate <95%. To perform the sex-mismatch analyses, variants on the X-chromosome were selected after filtering out low quality genotypes, call rate<95%, MAF<2%, low QD score (3 for INDELs and 2 for SNPs), low confidence regions and segmental duplications and those that do not have PASS VQSR. A set of high quality common autosomal variants were extracted for relative inference, principal component analysis, and estimation of heterozygosity after removing low confidence regions and segmental duplications, low quality genotypes, QD<2, call rate<98%, self-reported ancestry-specific HWE p>1×10^-6^ among controls, MAF<1% and do not have PASS VQSR. Heterozygosity was estimated within each ancestry and samples with F statistic>2 were removed. Genetically inferred ancestry was obtained using the 1000 Genomes as reference. Sample QC metrics were then calculated in HAIL using autosomal variants after filtering out low-quality genotypes, variants with ancestry-specific HWE p<1×10^-6^, low confidence regions and segmental duplications, low QD score (3 for INDELs and 2 for SNPs) and those that do not have PASS VQSR. Samples with call rate below 95%, mean DP below 30 and mean GQ below 80 were removed. Variant QC was done through filtering out monomorphic variants, call rate below 95%, those with HWE (p < 1 × 10^-6^), lie in low confidence regions or segmental duplications, are insertions or deletions with QD <= 3 or SNPs with QD <= 2 or variants where VQSR does not PASS unless singleton in which case retain those with VQSRTrancheSNP99.60to99.80. A total of 11,216 PC-identified European ancestry participants were included after additional removal of duplicates and relatives across data sources. A total of 2,734,519 variants were included.

The second UKB data set is a population-based dataset. Samples were filtered out if they showed mismatch between genetically determined and reported sex, high rates of heterozygosity or contamination (D-stat > 0.4), low sequence coverage (<85% of targeted bases achieving >20X coverage), duplicates, and exome sequence variants discordant with genotyping chip. More details are described elsewhere.^30^ The “Functionally Equivalent” (FE) call set was used.^31^ A total of 43,243 PC-identified European ancestry individuals were included after additional removal of duplicates and relatives across data sources.

### Variant Annotation

We compiled autosomal variants with call rate>95% within each case and ancestry specific analysis dataset with MAC≥1 (across the combined data). Variants were annotated using the Ensembl Variant Effect Predictor^32^ and its associated Loss-of-Function Transcript Effect Estimator (LOFTEE)^33^ and the dbNSFP^34^ version 3.5a plugins. We limited our annotations to the canonical transcripts. The LOFTEE plugin assesses stop-gained, frameshift, and splice site disrupting variants. Loss-of-function variants are classified as either high confidence or low confidence. The dbNSFP is a database that provides functional prediction data and scores for non-synonymous variants using multiple algorithms.^34^ This database was used to classify missense variants as damaging using two different definitions based on bioinformatic prediction algorithms. The first is based on MetaSVM^35^ which is derived from 10 different component scores (SIFT, PolyPhen-2 HDIV, PolyPhen-2 HVAR, GERP++, MutationTaster, Mutation Assessor, FATHMM, LRT, SiPhy, PhyloP). The second is based on 5 variant prediction algorithms including SIFT, PolyPhen-2 HumVar, PolyPhen-2 HumDiv, MutationTaster and LRT score. Additionally, we ran a deep neural network analysis (Splice AI) to predict splice-site altering variants.^36^ Multi-ancestry and ancestry-specific variant descriptive analyses were performed using variant-specific statistics obtained from the largest sample size out of the 6 phenotypes.

### Single-Variant Association Analysis

Each data source was sub-categorized based on ancestry and CAD or T2D case status in the studies ascertained by disease status. Subgrouping data sources yielded a total of 23 distinct sample sub-categories. As relatives were kept within each sub-group, we performed generalized linear mixed models to analyze the association of single autosomal variants with standard-deviation corrected-inverse-normal transformed traits using RVTESTS.^28^ RVTESTS was used to generate summary statistics and covariance matrices using 500 kilobase sliding windows. To obtain the single-variant associations, we performed a fixed-effects inverse-variance weighted meta-analysis for multi-ancestry and within each of the five major ancestries. An exome-wide significance threshold of P<7.2×10^-8^ (Bonferroni correction for six traits and using previously recommended threshold for coding variants P<4.3×10^-7^)^37^ was used to determine significant coding variants.

### Gene-Based Association Analysis

We used summary level score statistics and covariance matrices from autosomal single-variant association results to perform gene-based meta-analyses among all individuals and within each ancestry using RAREMETALS version 7.2.^38^ Samoan individuals only contributed to the overall analysis. Gene-based association testing aggregates variants within each gene unit using burden tests and SKAT which allows variable variant effect direction and size.^39^ The “rareMETALS.range.group” function was used with MAF<1%, which filters out all variants with combined MAF>1% in all meta-analytic datasets. All variants with call rates<95% and not annotated as LOF using LOFTEE, splice-site variants or damaging missense as defined by MetaSVM or by all SIFT, PolyPhen-2 HumVar, PolyPhen-2 HumDiv, MutationTaster and LRT prediction algorithms (Damaging 5 out of 5) were excluded in the gene-based meta-analyses.

We used 6 different variant groupings to determine the set of damaging variants within each gene, 1) high-confidence LOF using LOFTEE, 2) LOF and predicted splice-site altering variants, 3) LOF and MetaSVM missense variants, 4) LOF, MetaSVM missense and predicted splice-site altering variants, 5) LOF and damaging 5 out 5 missense variants, and 6) LOF, damaging 5 out 5 missense and predicted splice-site altering variants. An exome-wide significance threshold of P<4.3×10^-7^, Bonferroni corrected for the maximum number of annotated genes (n=19,540) and six lipid traits, was used to determine significant coding variants. Two gene transcripts, *DOCK6* and *DOCK7*, that overlap with two well-studied lipid genes, *ANGPTL8* and *ANGPTL3*, respectively, met our exome-wide significance threshold. After excluding variation observed in *ANGPTL8* and *ANGPTL3*, *DOCK6* and *DOCK7*, respectively, were no longer significant and have been excluded as associated genes.

We performed a series of sensitivity analyses for our results. We repeated the multi-ancestry gene-based analyses using a MAF<0.1%, and compared our exome-wide significant gene-based results using a MAF<1% to using a MAF<0.1%. We compared the single variants in our top gene-based associations with respective traits using GWAS summary data.^8^ Gene-based tests were repeated excluding variants identified in GWAS using P<5×10^-8^. Furthermore, all single variants included in each of the top gene-based association were analyzed in relation to the respective trait. To determine the variants that were contributing mostly to the gene-based signal, counts and proportions were obtained at different P value thresholds.

To understand whether variants contributing to top gene-based signals were similar or different across different ancestries, we determined the degree of overlap across ancestries for all variants incorporated and then for those with P<0.05. Finally, we checked for overlap across the most significant (lowest P value) variant from each of the gene-based signals.

Heterogeneity of gene-based estimates in all gene-trait-variant grouping combinations passing exome-wide significant levels was assessed across the five main ancestries (European, South Asian, African, Hispanic and East Asian) and between T2D and CAD cases and controls using Cochran’s Q.

### Replication of gene-based associations

We performed replication of our top gene-based associations with blood lipid levels in the Penn Medicine BioBank (PMBB) and UK Biobank samples that did not contribute to the discovery analysis.

The PMBB is a repository of genotype and phenotype data for 43,731 patients at the University of Pennsylvania Perelman School of Medicine. All individuals recruited for PMBB are patients of clinical practice sites of the University of Pennsylvania Health System. Appropriate consent was obtained from each participant regarding storage of biological specimens, genetic sequencing, and access to all available EHR data. The study was approved by the Institutional Review Board of the University of Pennsylvania and complied with the principles set out in the Declaration of Helsinki. The six lipid phenotypes studied were HDL-C (n=21,247), LDL-C (n=21,040), non-HDL-C (n=21,087), TC (n=21,153), TG (n=21,418), and TG:HDL (n=21,213). All available lipid trait measurements up to July 2020 were included. HDL-C, LDL-C, TC, and TG levels were measured directly and accessible via PMBB. Non-HDL-C levels were obtained by subtracting HDL-C from TC levels. TG and TG:HDL levels were logarithmically transformed to normalize their distribution for association testing. Due to the clinical nature of the biobank, samples often had multiple phenotype values corresponding to a patient’s various clinical appointments. Gene-based associations were performed on the minimum, median, and maximum phenotype values to account for both potentially protective and pathogenic effects. For the gene-based association analysis, 10 different variant groupings were used to determine the set of damaging variants within each gene including the six groupings used in the initial study. The additional four groupings used predicted loss-of-function (pLOF) variants that included frameshift, stop gain, and splicing variants as annotated by RefGene. Missense variants were annotated using Rare Exome Variant Ensemble Learner (REVEL) and filtered for those with a pathogenicity score>0.5. The four additional groupings consisted of, 1) pLOF, MAF≤0.1%, 2) pLOF, MAF≤0.1%, REVEL missense, 3) pLOF, MAF≤1%, and 4) pLOF, MAF≤1%, and REVEL missense. Each of the 10 groupings were used in a gene-based association test with the minimum, median, and maximum values of the 6 lipid phenotypes. Furthermore, ancestry-specific associations were also performed to elucidate any potential ancestry-specific effects. This included associations among African and European ancestries separately, and then the two populations meta-analyzed. All associations were adjusted for sex, age, and principal components. The first 5 principal components were used for African ancestry associations, and the first 10 principal components were used for European ancestry associations.

In UK Biobank, we analyzed the association of rare variant aggregates from the 10 genes against four lipid phenotypes in the UK biobank whole exome sequencing (WES) data. Variant aggregates were obtained for the following four categories 1) LOFTEE – HC 2) LOFTEE - HC & predicted splice site altering 3) LOFTEE - HC & deleterious-METAsvm 4) LOFTEE - HC & deleterious-METAsvm & predicted splice site altering. We removed UK Biobank individuals used in the discovery analysis, resulting in 150,694 individuals for replication. The phenotypes were adjusted for lipid lowering medications, where total cholesterol was adjusted by dividing by 0.8 and LDL-C by dividing by 0.7. Triglycerides were natural log transformed for analysis. The phenotypes were inverse rank normalized and scaled by the standard deviation of the trait and adjusted for covariates (sex, age, age2, PC1-PC10, if British ancestry). Rare variant aggregate test was conducted using STAAR^40^ with a MAF of 0.01 for the four lipids. Effect estimates were calculated using glmm.wald burden test. As a sensitivity analysis to determine the effects of statin treatment on the results, a similar analysis was carried out for a subset of individuals where samples with statins were removed, resulting in 127,459 individuals without statin treatment. For our top associations in our discovery, we found the effect sizes and p-value in the analysis including and excluding individuals on statin treatment remained similar (**Figure S1**).

### Gene-Based Analysis of GWAS Loci and Drug Targets

We performed gene-based analysis using the six variant groups for genes in GWAS loci. A locus was defined as the region around each GWAS index variant ± 200kb. Top GWAS signals were obtained from a recent meta-analysis of >300,000 individuals in the Million Veterans Program.^8^ In-silico lookup of gene-based associations for respective lipid traits were then performed for all genes within defined GWAS loci. Drug target genes were obtained from the drug bank database^41^ using the following search categories: “Hypolipidemic Agents, Lipid Regulating Agents, Anticholesteremic Agents, Lipid Modifying Agents and Hypercholesterolemia”. A liberal definition for drug targets was used – drugs with any number of targets and targets targeted by any number of drugs – and then in-silico lookups were performed for gene-based associations.

### Gene-set Enrichment Analysis

Gene-set enrichment analyses were performed for sets of Mendelian-, protein-altering- and non-protein altering GWAS, and drug target genes with LDL-C, HDL-C and TG. 21 Mendelian genes were included based on previous literature^2^: *LDLR*, *APOB*, *PCSK9*, *LDLRAP1*, *ABCG5*, *ABCG8*, *CETP*, *LIPC*, *LIPG*, *APOC3*, *ABCA1*, *APOA1*, *LCAT*, *APOA5*, *APOE*, *LPL*, *APOC2*, *GPIHBP1*, *LMF1*, *ANGPTL3*, and *ANGPTL4*. We analyzed GWAS gene sets based on their coding status and their proximity to the most significant signal in the GWAS. Coding variants were defined as missense, frameshift, or stop gained variants. Gene sets for coding or non-coding variants were then stratified into three categories based on proximity to the most significant variant within each locus – closest-, second closest- and greater than second closest gene. For each gene within each set, we obtained the most significant association in the multi-ancestry or ancestry specific meta-analysis set using any of the six different variant groups. Then each gene within each gene set was matched to 10 other genes based on sample size, total number of variants, cumulative MAC, and variant grouping nearest neighbors using the matchit R function. Then we compared the proportions using Fisher’s exact test between the main and matched gene sets by applying different P-value thresholds.

### Association of Lipid Genes with CAD and T2D data and liver fat/markers

We determined the associations of 40 genes identified in the main and GWAS loci analyses with CAD, T2D, and glycemic and liver enzyme blood measurements. The association with T2D was obtained from the latest gene-based exome association data from the AMP-T2D-GENES consortium.^15^ The reported associations were obtained from different variant groups based on their previous analyses. We additionally performed gene-based association analyses with CAD using the MIGen case-control, UKB case-control, and UKB cohort samples using the variant groups described above. Further, six traits including fasting plasma glucose, HbA1c, alanine aminotransferase, aspartate aminotransferase, gamma glutamyl transferase and albumin were analyzed in the UKB dataset. Single variant association analyses were performed with RVTESTS. Linear mixed models incorporating kinship matrices were used to adjust for relatedness within each study. Covariance matrices were generated using 500 kilobase sliding windows. RAREMETALS was used to assess associations between aggregated variants (MAF<1%) in burden and SKAT tests with CAD and each of the six quantitative traits. We used 6 different variant groupings to determine the set of damaging variants within each gene, 1) high-confidence LOF using LOFTEE, 2) LOF and predicted splice-site altering variants, 3) LOF and MetaSVM missense variants, 4) LOF, MetaSVM missense and predicted splice-site altering variants, 5) LOF and damaging 5 out 5 missense variants, and 6) LOF, damaging 5 out 5 missense and predicted splice-site altering variants.

## Results

### Sample and variant characteristics

Individual-level, quality-controlled data were obtained from four sequenced study sources with circulating lipid data for individuals of multiple ancestries (Figure 1). Characteristics of the study samples are detailed in **Table S1**. We analyzed data on up to 172,000 individuals with LDL-C, non-HDL-C (a calculated measure of TC minus HDL-C), TC, HDL-C, TG, and TG:HDL ratio (a proxy for insulin resistance).^42; 43^ 56.7% (n=97,493) of the sample are of European ancestry, 17.4% (n=30,025) South Asian, 9.6% (n=16,507) African American, 9.6% (n=16,440) Hispanic, 6.1% (n=10,420) East Asian, and 0.7% (n=1,182) Samoan, based on genetically-estimated and/or self-reported ancestry.

**Figure 1.**
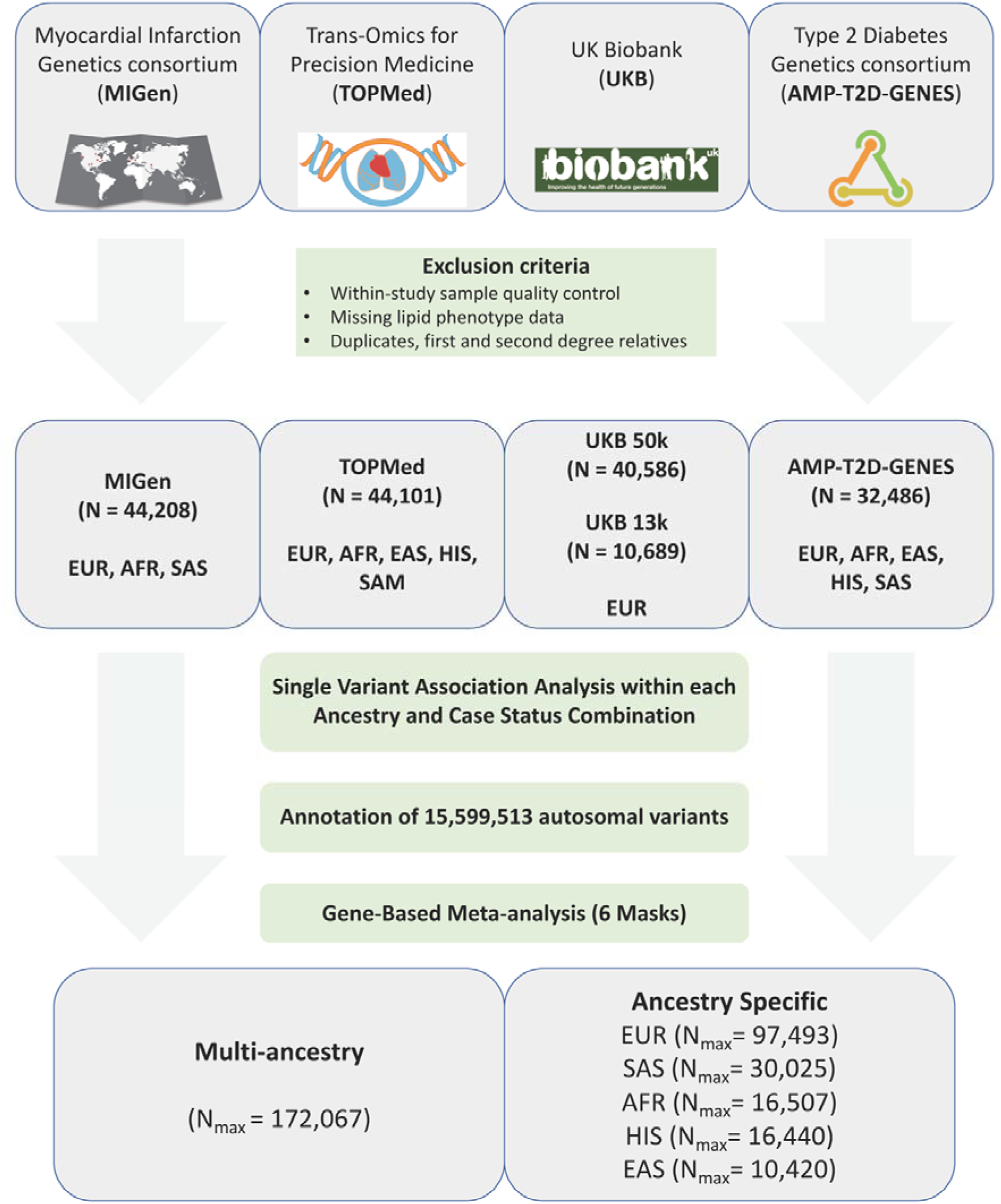
Study samples and design. Flow chart of the different stages of the study. Exome sequence genotypes were derived from four major data sources: The Myocardial Infarction Genetics consortium (MIGen), the Trans-Omics from Precision Medicine (TOPMed), the UK Biobank and the Type 2 Diabetes Genetics (AMP-T2D-GENES) consortium. Single-variant association analyses were performed by ancestry and case-status in case-control studies and meta-analyzed. Single-variant summary estimates and covariance matrices were used in gene-based analyses using 6 different variant groups and in multi-ancestry and each of the five main ancestries. AFR=African ancestry, EAS=East Asian ancestry, EUR=European ancestry, HIS=Hispanic ancestry, SAM=Samoan ancestry, SAS=South Asian ancestry

After sequencing, we observed 15.6 M variants across all studies; 5.0 M (32.6%) we classified as transcript-altering coding variants based on an annotation of frameshift, missense, nonsense, or splice site acceptor/donor using the Variant Effect Predictor (VEP).^32^ A total of 340,214 (6.7%) of the coding variants were annotated as high confidence loss-of-function (LOF) using the LOFTEE VEP plugin,^33^ 238,646 (4.7%) as splice site altering identified by Splice AI,^36^ 729,098 (14.3%) as damaging missense as predicted by the MetaSVM algorithm^35^, and 1,106,309 (21.8%) as damaging missense as predicted by consensus in all five prediction algorithms (SIFT, PolyPhen-2 HumVar, PolyPhen-2 HumDiv, MutationTaster and LRT).^34^ As expected, we observed a trend of decreasing proportions of putatively deleterious variants with increasing allele count (**Figure S2, Table S3**).

### Single-variant association

We performed inverse-variance weighted fixed-effects meta-analyses of single-variant association results of LDL-C, non-HDL-C, TC, HDL-C, TG and TG:HDL ratio from each consortium and ancestry group. Meta-analysis results were well controlled with genomic inflation factors ranging between 1.01 and 1.04 (**Table S4**). Single-variant results were limited to the 425,912 protein-altering coding variants with a total minor allele count (MAC) > 20 across all 172,000 individuals. We defined significant associations by a previously established exome-wide significance threshold for coding variants (P<4.3×10^-7^)^37^ which was additionally corrected for testing six traits (P=4.3×10^-7^ divided by 6) within all study samples or within each of the five major ancestries (**Tables S5-S10**); this yielded in each analysis a significance threshold of P<7.2×10^-8^. A total of 104 rare coding variants in 57 genes were associated with LDL-C, 95 in 54 genes with non-HDL-C, 109 in 65 genes with TC, 92 in 56 genes with HDL-C, 61 in 36 genes with TG, and 68 in 42 genes with TG:HDL. We identified six missense variants in six genes (*TRIM5* p.Val112Phe, *ADH1B* p.His48Arg, *CHUK* p.Val268Ile, *ERLIN1* p.Ile291Val, *TMEM136* p.Gly77Asp, *PPARA* p.Val227Ala) >1Mb away from any index variant previously associated with a lipid phenotype (LDL-C, HDL-C, TC, or TG) in previous genetic discovery efforts (**Tables S5-S10**).^3; 7; 8^ *PPARA* p.Val227Ala has previously been associated with blood lipids at a nominal significance level in East Asians (P < 0.05), where it is more common than in other ancestries.^44^ Both *TRIM5* and *ADH1B* LDL-C increasing alleles have been associated with higher risk of CAD in a recent GWAS from CARDIOGRAM (OR: 1.08, P=2×10^-9^; OR=1.08, P=4×10^-4^).^45^ Single variant associations were further performed in each of the five main ancestries (**Table S11**).

### Gene-based association

Next we performed gene-based testing of transcript-altering variants in aggregated burden and sequence kernel association tests (SKAT)^46^ tests in all study participants and within each of the six main ancestries for six lipid traits: LDL-C, HDL-C, non-HDL-C, TC, TG, and TG:HDL. We excluded the Samoans from the single-ancestry analysis given the small number of individuals. We limited attention to variants with MAF≤1% for each of six variant groups: 1) LOF, 2) LOF and predicted splice-site altering variants using Splice AI, 3) LOF and MetaSVM missense variants, 4) LOF, MetaSVM missense and predicted splice-site altering variants, 5) LOF and damaging 5 out 5 missense variants, and 6) LOF, damaging 5 out 5 missense and predicted splice-site altering variants. Meta-analyses results were well controlled (**Table S12**).

We identified 35 genes reaching exome-wide significance (P=4.3×10^-7^) for at least one of the six variant groupings (**Tables S13-S19**). Most of the significant results were from the multi-ancestry analysis, with multiple ancestries contributing to the top signals (Figure 2A) and most of the 35 genes were associated with more than one lipid phenotype (Figure 2B). Ten of the 35 genes did not have prior evidence of gene-based links with blood lipid phenotypes (**Table 1**), and seven genes, including *ALB*, *SRSF2*, *CREB3L3*, *NR1H3*, *PLA2G12A*, *PPARG*, and *STAB1* have evidence for a biological connection to circulating lipid levels (Box 1).

#### Box 1. Genes with biological links to lipid metabolism

|  |  |
| --- | --- |
| **ALB** | The association between mutations in the albumin gene and elevated cholesterol levels has been previously observed in rare cases of congenital analbuminemia.<sup>54</sup> This has been mainly suggested to result from compensatory increases in hepatic production of other non-albumin plasma proteins to maintain colloid osmotic pressure particularly apolipoprotein B-100 leading to elevations in TC and LDL-C but normal HDL-C levels – which is consistent with our findings – although the exact mechanisms remain uncertain.<sup>55</sup> A lipodystrophy-like phenotype has also been linked to analbuminemia which is consistent with the suggestive tendency for increased risk of T2D with LOF and predicted damaging variants in albumin in the population (OR=1.85; P=0.007) (**Supplementary Table 30**). |
| **SRSF2** | The *SRSF2* gene encodes a highly conserved serine/arginine-rich splicing factor and has previously been linked to acute liver failure in liver-specific knockout in mice with accumulation of TC in the mutant liver.<sup>56</sup> Thus, this gene could be linked to a non-alcoholic fatty liver phenotype with accumulation of lipids in the liver as observed with other genes as *PNPLA3* and *TM6SF2*.<sup>7</sup> Therefore, we looked at association with liver function markers and we found an association between *SRSF2* and higher albumin levels ($P = 1 \times 10^{-4}$) and a suggestive tendency for higher gamma glutamyl transferase (GGT) ($P = 0.05$), consistent with potential liver involvement (**Supplementary Table 33-36**). |
| **CREB3L3** | The association between *CREB3L3* and higher TG supports previous evidence from a single family and cohorts with severe hypertriglyceridemia but not sufficient evidence to be classified as a Mendelian lipid gene (ref).<sup>57-59</sup> This has been additionally supported by functional studies where *Creb3l3* knockout mice showed hypertriglyceridemia partly due to deficient expression of lipoprotein lipase coactivators (*Apoc2*, *Apoa4*, and *Apoa5*) and increased expression of activator *Apoc3*.<sup>58</sup> |
| **NR1H3** | The observed association of *NR1H3* with higher HDL-C and lower TG is supported by previous evidence of a role in non-alcoholic fatty liver disease in mice.<sup>60</sup> This gene encodes a liver X receptor alpha (LXRα) which is a nuclear receptor that acts as a cholesterol sensor and protects from cholesterol overload.<sup>61; 62</sup> It has previously been shown that disrupting the LXRα phosphorylation at Ser196 in mice prevents non-alcoholic fatty liver disease.<sup>60</sup> |
| **PLA2G12A** | *PLA2G12A* is in the secretory phospholipase A2 (sPLA<sub>2</sub>) family, which liberates fatty acids in the -sn2 position of phospholipids. This pattern suggests a previously unreported possible lipolytic role of this phospholipase in a manner similar to another member of the adipose-specific phospholipases, *PLA2G16*, which has been shown to have a lipolytic role in mice.<sup>63; 64</sup> Further studies are needed to confirm whether *PLA2G12A* has a lipolytic role. |
| **PPARG** | Rare loss of function mutations in *PPARG* have been previously found to be associated with reduced adipocyte differentiation, lipodystrophy and increased risk of T2D.<sup>65-67</sup> |
| **STAB1** | The *STAB1* gene is a scavenger receptor that has been shown to mediate uptake of oxidized LDL-C.<sup>68; 69</sup> There was a suggestive association between LOF variants and higher LDL-C ($\beta = 4.3$ mg/dL, $P = 2 \times 10^{-3}$) consistent with its role in LDL-C uptake. |

**Figure 2.**
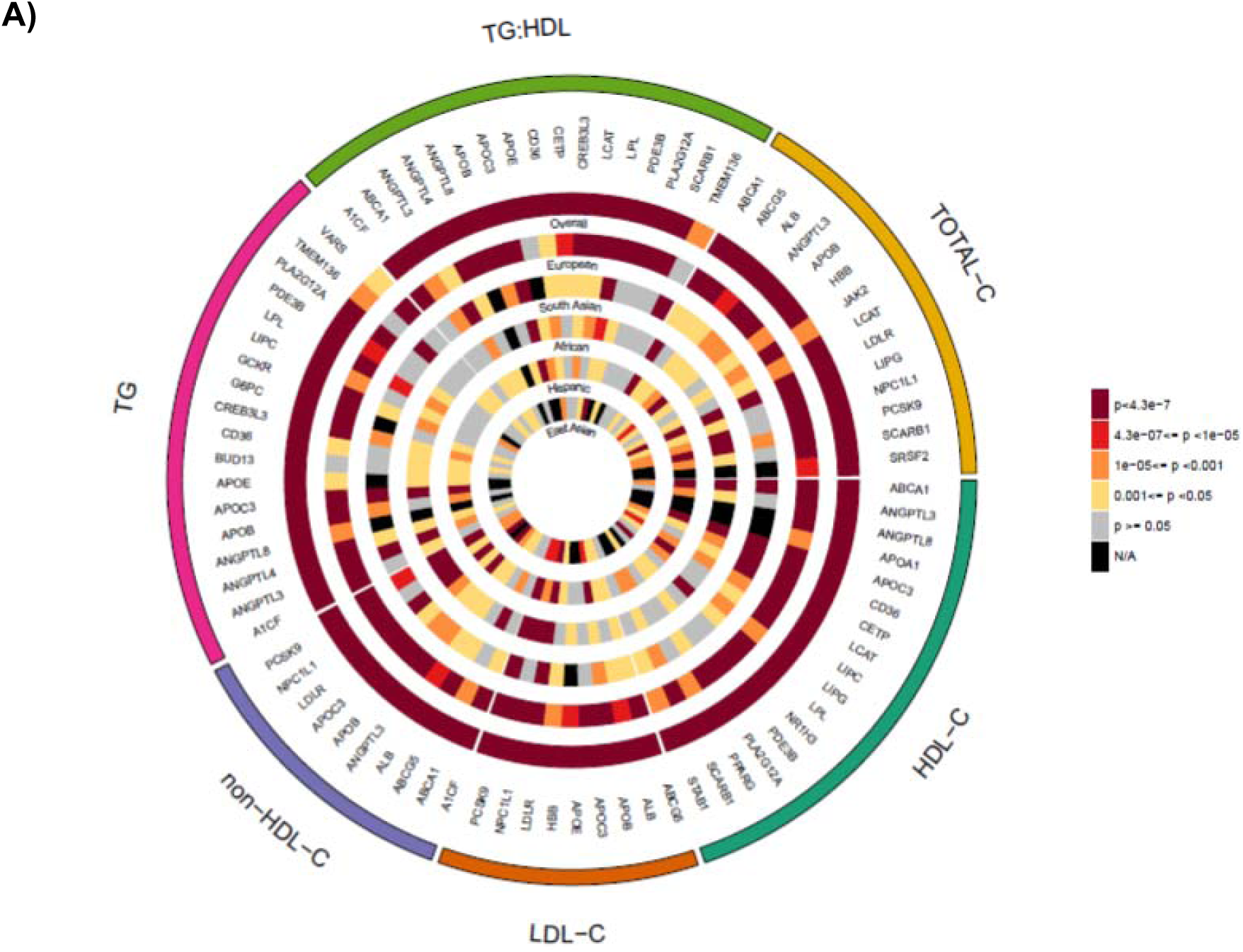
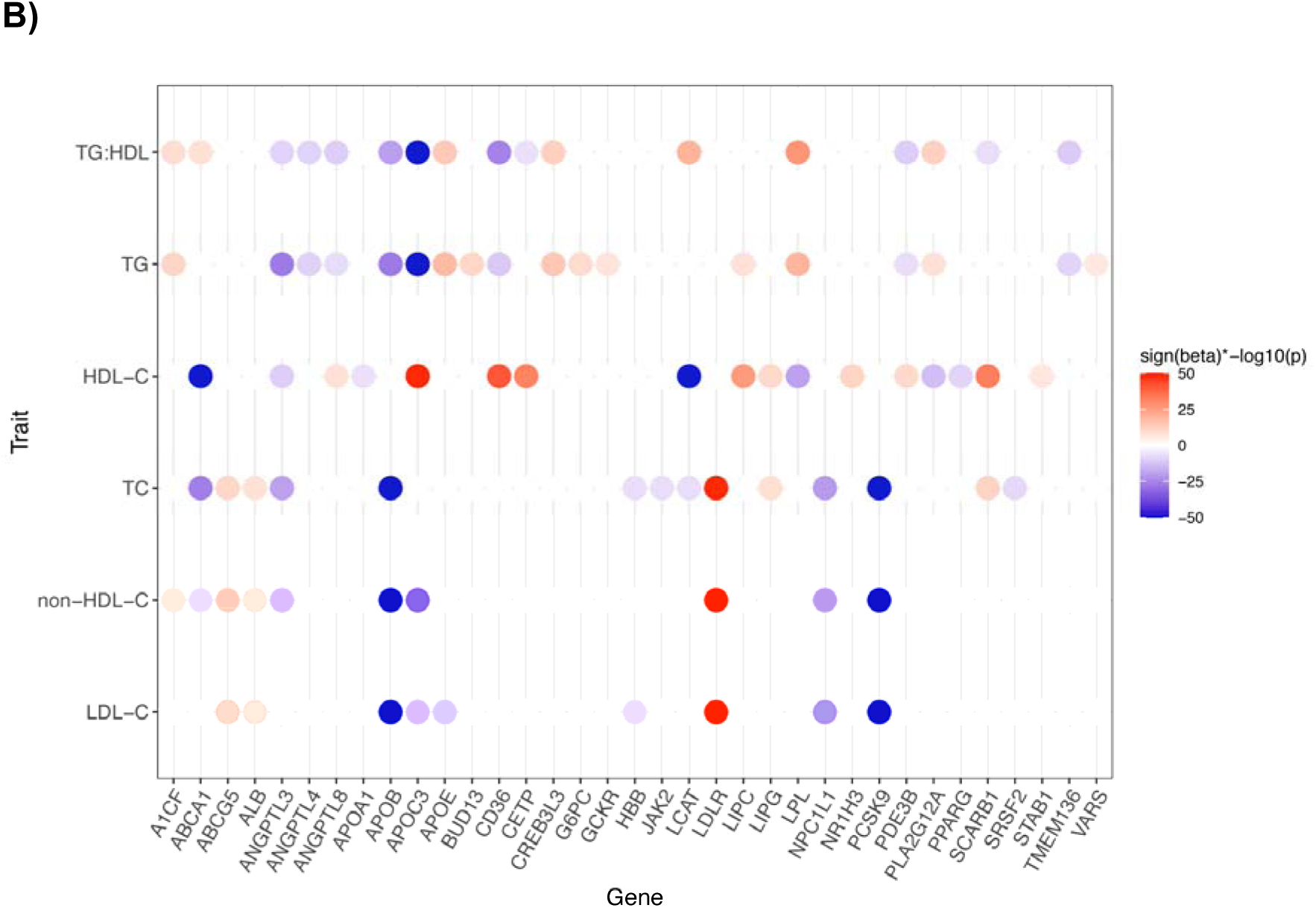
Exome-wide significant associations with blood lipid phenotypes. **A)** Circular plot highlighting the evidence of association between the exome-wide significant 35 genes with any of the six different lipid traits (P < 4.3 × 10^-7^). The most significant associations from any of the six different variant groups are plotted. For almost all of the genes the most significant associations were obtained from the multi-ancestry meta-analysis. **B)** Strength of association of the 35 exome-wide significant genes based on the most significant variant grouping and ancestry across the six lipid phenotypes studied. Most of the genes indicated associations with more than one phenotype. Sign(beta)*-log10(p) displayed for associations that reached a P < 4.3 × 10^-7^. When the Sign(beta)*-log10(p) > 50, they were trimmed to 50.

We performed a series of sensitivity analyses on our results. To determine whether low frequency variants between 0.1%-1% frequency were driving our gene-based association results, we performed the gene-based multi-ancestry meta-analyses using a maximum MAF threshold of 0.1% instead of 1%. We observed exome-wide significant associations (P<4.3×10^-7^) for 29 genes using a 0.1% MAF threshold, all observed in our primary analyses using a MAF threshold of 1% (**Table S20**). We then intersected our 35 lipid associated genes from 85 gene-based associations observed in the primary analysis with our results using a MAF threshold of 0.1%. All genes remained at least nominally significant (P < 0.05) using a 0.1% MAF threshold, except the *A1CF* and *TMEM136* associations (**Table S21**). Furthermore, we determined whether those signals were driven by previously reported GWAS hits. We identified a total of 7 HDL-C associated variants in 6 genes, 7 LDL-C variants in 3 genes, 3 TC variants in 1 gene and 7 TG variants in 6 genes that were previously found to be genome-wide significant in MVP (**Table S22**).^8^ Respective gene-based analyses were repeated without those variants. Gene-based signals at *A1CF* and *BUD13* were lost after removal of 1 variant in each of those genes (**Table S23**).

The *JAK2* signal was further investigated after splitting the 136 contributing variants into those annotated as somatic using the Catalogue Of Somatic Mutations In Cancer (COSMIC) database and not annotated as a somatic variant. We observed an association only among a set of 26 variants annotated as somatic while no association was observed using the remaining 110 variants (**Table S24**).

We also determined which of the 35 genes were outside GWAS regions defined as those within ±200kb flanking regions of GWAS indexed Single nucleotide polymorphisms (SNPs) for TC (487 SNPs), LDL-C (531 SNPs), HDL-C, and TG (471 SNPs).^8^ We identified 1,295 unique genes included in these lipid GWAS regions. Eight out of the 35 associated genes (23%) were not within a GWAS region (**Table S13**).

To understand whether the gene-based signals were driven by variants that could be identified through single variant analyses, we looked at the proportion of the 35 genes that were associated with each trait that have at least one single contributing variant that passed the genome-wide significance threshold of 5×10^-8^. Seventeen genes were associated with HDL-C at exome-wide significance (**Table S13**); eight genes had at least one variant with P<5×10^-8^ (**Table S8**). Similarly, we observed 4/9 for LDL-C, 4/10 non-HDL-C, 4/14 TC, 7/18 TG, and 6/17 TG:HDL genes with at least one genome-wide significant variant (**Tables S5-S10**).

For genes with both gene-based and single variant signals, we determined the variants were driving these signals, and determined the single variant associations for all variants contributing to the top 35 genes (**Table S25**). From a total of 85 gene-based associations, 33 had at least one and 19 had only one single variant with P<5×10^-8^ (**Tables S25 and S26**). All of the 19 had at least 2 variants passing nominal significance (P<0.05) and 13 had at least 10 variants with P <0.05.

### Comparison of gene-based associations across ancestries

We determined the overlap between single variants included in gene-based signals across the five main ancestries. A large proportion of variants from each ancestry did not overlap with any other ancestry (**Figure S3**). For example, a total of 15 genes were observed to have significant gene-based associations with HDL-C in multi-ancestry meta-analyses. A total 69% of variants from European ancestry samples that contributed to HDL-C gene-based associations did not overlap with any other ancestry, and was 60% in South Asian-, 44% in African-, 40% in Hispanic- and 59% in East Asian ancestry. When restricted to variants with P<0.05 in the multi-ancestry meta-analysis, the overlap among ancestries increased (**Figure S4**). A total 57% of variants from European ancestry did not overlap with any other ancestry, and was 42% in South Asian-, 20% in African-, 24% in Hispanic- and 34% in East Asian ancestry. Finally, we determined the top single variant contributing to each gene-based association (**Figure S5**). None of the top variants overlapped with all ancestries and 80% of EUR variants did not overlap with any other ancestry, and was 87% in South Asian-, 93% in African-, 80% in Hispanic- and 93% in East Asian ancestry.

But, the gene-based associations were mostly consistent across the six ancestry groupings: European, South Asian, African, Hispanic, and East Asian. Three of the 17 HDL-C genes showed association in at least two different ancestries at exome-wide significance level (P=4.3×10^-7^). Similarly, 3/9 LDL-C, 4/10 non-HDL-C, 5/14 TC, 2/18 TG and 2/17 TG:HDL genes showed association in at least two difference ancestries at a exome-wide significance level. Using a less stringent significance level (P<0.01), across the six lipid traits, 59-89% of associated genes from the joint analysis were associated in at least two different ancestries.

We tested the top 35 genes for heterogeneity across all 303 gene-trait-variant grouping combinations passing the exome-wide significance threshold (P<4.3×10^-7^). We observed heterogeneity in effect estimates (P_Het_<1.7×10^-4^, accounting for 303 combinations) in 19 (6%) different gene-trait-variant grouping combinations and in six different genes: *LIPC*, *LPL*, *LCAT*, *ANGPTL3*, *APOB*, and *LDLR* (**Table S27**). Although the LOF gene-based effect sizes were largely consistent across ancestries, there were differences in the cumulative frequencies of LOF variants for several genes including *PCSK9*, *NPC1L1*, *HBB* and *ABCG5* (**Figures S6-S8**).

We observed LOF and predicted damaging variants in the *TMEM136* gene associated with TG and TG:HDL only among individuals of South Asian ancestry (P_SKAT_=3×10^-9^ and 2×10^-11^, respectively) (**Table 1**, Figure 2A). With the same variant grouping and ancestry, we observed associations with reduced TG by burden tests (β =-15%, P=3×10^-4^) and TG:HDL (β =-20%, P=6×10) (**Tables S18 and S19**). Additionally, a single missense variant was associated only among South Asians (rs760568794,11:120327605-G/A, p.Gly77Asp) with TG (β =-36.9%, P=2×10^-8^) (**Table S9**). This variant was present only among South Asian (MAC=24) and Hispanics (MAC=8), but showed no association among Hispanics (P=0.86). This gene encodes a transmembrane protein of unknown function.

### Replication of gene-based associations

We performed replication using the Penn Medicine BioBank (PMBB) and UK Biobank samples that did not contribute to the initial analysis. In PMBB, we observed associations at a nominal significance level (p<0.05) and in the same direction as the discovery for 6 out of the 10 genes without prior evidence of gene-based links with blood lipid phenotypes (*SRSF2, JAK2, CREB3L3, NR1H3, PLA2G12A, PPARG*) with their respective blood lipids. For the gene *TMEM136*, we found an association of nominal significance for TG and TG:HDL as well, but with a beta in the opposite and positive direction. For the other 3 genes, *ALB, VARS, and STAB1*, we did not find associations at a nominal significance level for their respective blood lipid traits (**Table S28**). In UK Biobank, we found 7 of the 10 genes were associated at a nominal significance level and in the same direction of effect as the discovery analysis (ALB*, JAK2, CREB3L3, NR1H3, PLA2G12A, PPARG, STAB1*) (**Table S29**). The only two genes that did not show evidence of replication in at least one of the replication studies were *TMEM136* and *VARS*. This may indicate these associations are false positives or that we lack power for replication for these associations. Our replication studies did not include individuals of South Asian ancestry and we observed that our association of *TMEM136* with TG and TG:HDL is driven by individuals of South Asian ancestry.

### Comparison of gene-based associations by case-status

We analyzed heterogeneity by CAD or T2D case status for the top 35 genes. The top 85 signals presented in **Table S13** determined in case-status specific meta-analyses for CAD and T2D. Out of the 85 different gene-based associations, we observed minimal heterogeneity in the results by case status. *LDLR*, *LCAT* and *LPL* showed significant heterogeneity by CAD case status and *LCAT* and *ANGPTL4* by T2D status (P_Het_ < 6×10^-4^) (**Tables S30 and S31**).

### Gene-based associations in GWAS loci

We determined whether genes near lipid array-based GWAS signals^8^ were associated with the corresponding lipid measure using gene-based tests of rare variants with the same traits. We obtained genes from 200 Kb flanking regions on both sides of each GWAS signal; 487 annotated to LDL-C GWAS signals, 531 to HDL-C signals, and 471 to TG signals. We analyzed genes within these three sets for gene-based associations with their associated traits. A total of 13, 19, and 13 genes were associated (P<3.4×10^-5^, corrected for the number of genes tested for the three traits) with LDL-C, HDL-C or TG, with 32 unique genes identified in the GWAS loci (**Tables S32-S37)**.

Three of the 32 genes had no prior aggregate rare variant evidence of blood lipid association. Variants annotated as LOF or predicted damaging in *EVI5* were associated with LDL-C (P_SKAT_=2×10^-5^). The burden test showed association with higher LDL-C levels (β=1.9 mg/dL, P=0.008) (**Table S32)**. Variants annotated as LOF or predicted damaging in *SH2B3* were associated with lower HDL-C (1=-2.5 mg/dL, P=1×10^-6^) among Europeans and variants that were annotated as LOF in *PLIN1* were associated with higher HDL-C (β= 3.9 mg/dL, P=1×10) (**Table S33**). Other genes in the regions of *EVI5*, *SH2B3*, and *PLIN1* did not show an association with the corresponding lipid traits (P>0.05) in multi-ancestry analyses. A previous report implicated two heterozygous frameshift mutations in *PLIN1* in three families with partial lipodystrophy.^47^ The gene encodes perilipin, the most abundant protein that coats adipocyte lipid droplets and is critical for optimal TG storage.^48^ We observed a nominal associations of *PLIN1* with TG (β =-7.0%, P=0.02). Our finding is contrary to what would be expected with hypertriglyceridemia in a lipodystrophy phenotype given the association with lower TG. This gene has an additional role where silencing in cow adipocytes has been shown to inhibit TG synthesis and promote lipolysis,^49^ which may explain those contradictions.

### Enrichment of Mendelian-, GWAS-, and drug targets genes

We next sought to test the utility of genes that showed some evidence for association but did not reach exome-wide significance. Within the genes that reached a sub-threshold level of significant association in this study using burden or SKAT tests (p < 0.005), we determined the enrichment of i) Mendelian dyslipidemia (N=21 genes)-;^2^ ii) lipid GWAS (N=487 for LDL-C, N=531 for HDL-C and N=471 for TG)^8^; and iii) drug target genes (N=53).^41^ We stratified genes in GWAS loci according to coding status of the index SNP and proximity to the index SNP (nearest gene, second nearest gene, and genes further away). We tested for enrichment of gene-based signals (P<0.005) in the gene sets compared to matched genes (Figure 3). For each gene within each gene set, the most significant association in the multi-ancestry or an ancestry specific analysis was obtained and then matched to 10 genes based on sample size, total number of variants, cumulative MAC, and variant grouping. The strongest enrichment was observed for Mendelian dyslipidemia genes within the genes that reached P < 0.005 in our study. For example, 52% of the HDL-C Mendelian genes versus 1.4% of the matched set reached P < 0.005 (OR:71, 95% CI: 16-455). We also observed that 45.5% of the set of genes closest to an HDL-C protein-altering GWAS variant reached P < 0.005 versus 1.4% in the matched gene set (OR:57, 95% CI: 13-362). Results were significant but much less striking for genes at non-coding index variants. We observed that 8.9% of the set of genes closest to an HDL-C non-protein altering GWAS variant reached P < 0.005 versus 2.3% in the matched set (OR:4.1, 95% CI: 1.8-8.7). While 8% of the set of genes in the second closest to an HDL-C non-protein altering GWAS variant reached P <0.005 versus 2.6% in the matched set (OR: 3, 95% CI: 1.1-8.3). There was no significant enrichment in second closest or >= third closest genes to protein altering GWAS signals and in >= third closest genes to non-protein altering GWAS signals. Drug target genes were significantly enriched in LDL-C gene-based associations (OR: 5.3, 95% CI: 1.4-17.8) but not in TG (OR: 2.2, 95% CI: 0.2-11.2) or HDL-C (OR: 1.0, 95% CI: 0.1-4.3) (Figure 3 **and Tables S38-S41**).

**Figure 3.**
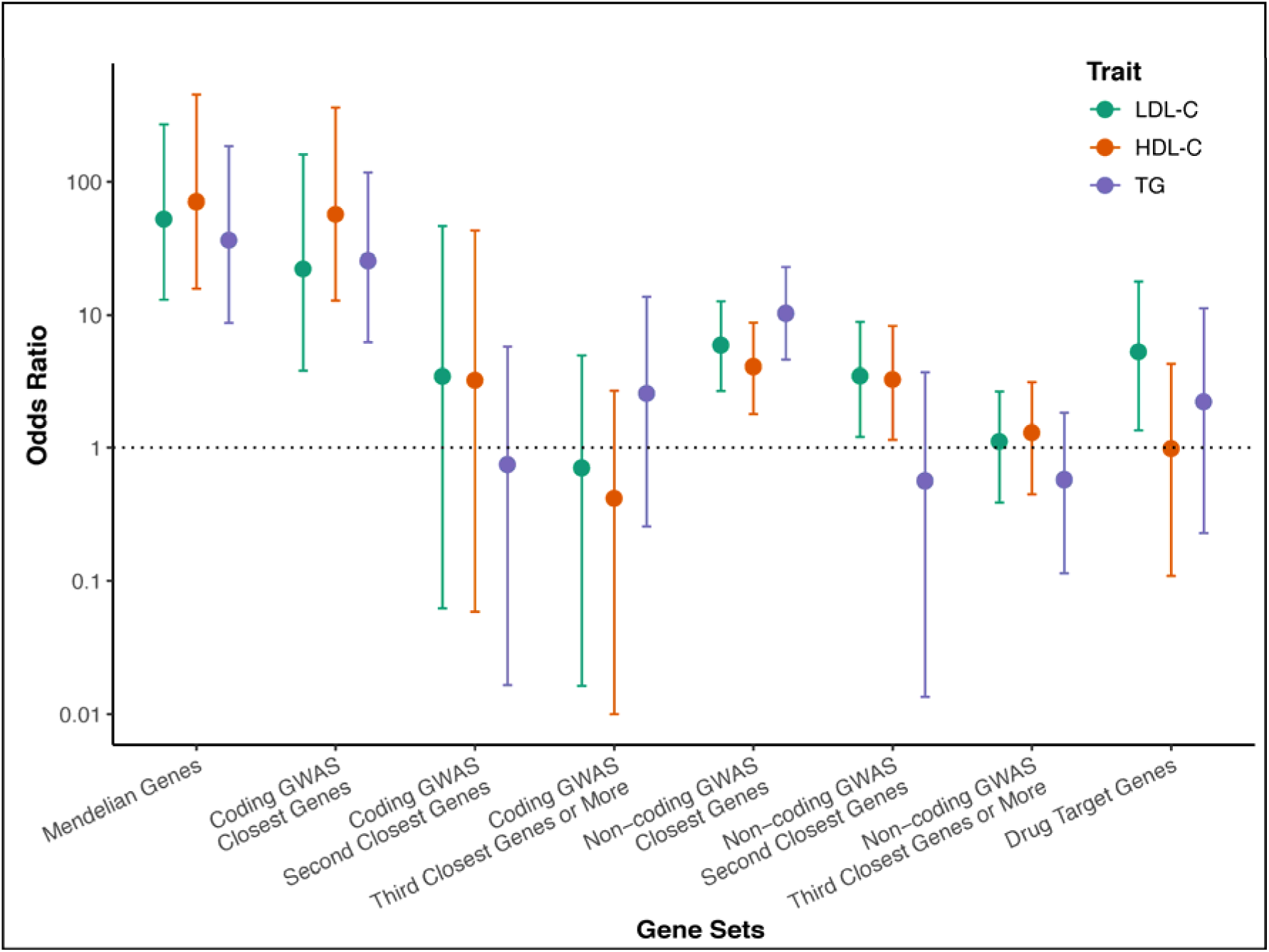
Enrichment of Mendelian, GWAS, and drug target genes in the gene-based lipid associations. Enrichment of gene sets of Mendelian genes (n=21), GWAS loci for LDL-C (n=487), HDL-C (n=531), and triglycerides (TG) (n=471) genes and drug target genes (n=53).

### Association of lipid genes with CAD, T2D, glycemic traits, and liver enzymes

We tested the genes identified through our main (35 genes) and GWAS loci (32 genes) for associations with CAD or T2D in our gene-based analyses (40 genes across the two sets). The CAD analyses were restricted to a subset of the overall exome sequence data with information on CAD status which included the MIGen CAD case-control, UK Biobank (UKB) CAD nested case-control, and the UKB cohort with a total of 32,981 cases and 79,879 controls. We observed four genes significantly associated with CAD (P_CAD_<0.00125, corrected for 40 genes). The four genes associated with lipids and CAD were all primarily associated with LDL-C: *LDLR* (OR: 2.97, P=7×10^-24^), *APOB* (P_SKAT_=4×10^-5^), *PCSK9* (OR: 0.5, P=2×10^-4^) and *JAK2* (P_SKAT_=0.001). Several other known CAD associated genes (*NPC1L1*, *CETP*, *APOC3*, and *LPL*) showed nominal significance for association with lipids (P<0.05). We observed nominal associations with CAD for two of the newly-identified lipid genes: *PLIN1* (P_SKAT_=0.002) and *EVI5* (OR: 1.29, P=0.002; **Table S42**). None of the 40 lipid genes reached significance for association with T2D in the latest AMP-T2D exome sequence results. We observed nominal associations of T2D with *STAB1* (OR: 1.05, P_T2D_=0.002) and *APOB* (OR: 1.08, P_T2D_=0.005) (**Table S43**).^15^

We additionally tested the 40 genes for association with six glycemic and liver biomarkers in the UKB: blood glucose, HbA1c, alanine aminotransferase (ALT), aspartate aminotransferase (AST), gamma glutamyl transferase (GGT), and albumin (**Tables S44-S49**). Using an exome-wide significance threshold of P=0.0012, we found associations between *PDE3B* and elevated blood glucose, *JAK2* and *SH2B3* and lower HbA1c, and *APOC3* and higher HbA1c. We found associations between *CREB3L3* and lower ALT, ALB, and higher AST, and between *A1CF* and higher GGT. *ALB* and *SRSF2* were associated with lower and higher albumin levels, respectively (**Tables S44-S49**).

## Discussion

We conducted a large multi-ethnic study to identify genes in which protein-altering variants demonstrated association with blood lipid levels. First, we confirm previous associations of genes with blood lipid levels and show that we detect associations across multiple ancestries. Second, we identified gene-based associations that were not observed previously. Third, we show that along with Mendelian lipid genes, the genes closest to both protein altering and non-protein altering GWAS signals, and LDL-C drug target genes have the highest enrichment of gene-based associations. Fourth, of the new gene-based lipid associations, *PLIN1* and *EVI5* showed suggestive evidence of an association with CAD.

Our study found that evidence of gene-based associations for the same gene in multiple ancestries. The heterogeneity in genetic association with common traits and complex diseases has been discussed extensively. A recent study has shown significant heterogeneity across different ancestries in the effect sizes of multiple GWAS identified variants.^50^ However, our study shows that gene-based signals are detected in multiple ancestries with limited heterogeneity in the effect sizes. Our study highlights enrichment of gene-based associations for Mendelian dyslipidemia genes, genes with protein-altering variants identified by GWAS, and genes that are closest to non-protein altering GWAS index variants. A previous transcriptome-wide Mendelian randomization study of eQTL variants indicated that most of the genes closest to top GWAS signals (>71%) do not show significant association with the respective phenotype.^51^ In contrast, our study provides evidence from sequence data that the closest gene to each top non-coding GWAS signal is most likely to be the causal one, indicating an allelic series in associated loci. This has implications for GWAS results, suggesting the prioritization of the closest genes for follow-up studies. We also observed enrichment of drug target genes only among LDL-C gene-based associations and not for HDL-C and TG gene-based associations, consistent with the fact that most approved therapeutics for cardiovascular disease targeting LDL-C

The gene-based analyses of lipid genes with CAD confirmed previously reported and known associations (*LDLR*, *APOB*, and *PCSK9).* Using a nominal P threshold of 0.05 we also confirmed associations with *NPC1L1*, *CETP*, *APOC3*, and *LPL*. Of the novel lipid genes, we observed borderline significant signals with *EVI5* and higher risk of CAD and between *PLIN1* and lower risk of CAD. The putative cardio-protective role of PLIN1 deficiency is supported by previous evidence in mice which has indicated reduced atherosclerotic lesions with Plin1 deficiency in bone marrow derived cells.^52^ This suggests PLIN1 as a putative target for CAD prevention; however, replication of the CAD association would be needed to confirm those signals.

There are limitations to our results. First, we had lower sample sizes for the non-European ancestries, limiting our power to detect ancestry-specific associations, and detect replication for *TMEM136* that was driven by a variant in South Asians. However, we find consistency of results across ancestries, and when we relax our significance threshold, the majority of associations (59-89%) are observed in more than one ancestry. Second, it has been reported that there was an issue with the UKB functionally equivalent WES calling.^53^ This mapping issue may have resulted in under-calling alternative alleles and therefore should not increase false positive findings. Third, we relied on a meta-analysis approach using summary statistics to perform our gene-based testing due to differences in sequencing platforms and genotyping calling within the multiple consortia contributing to the results. This approach has been shown to be equivalent to a pooled approach for continuous outcomes.^38^

In summary, we demonstrated association between rare protein-altering variants with circulating lipid levels in >170,000 individuals of diverse ancestries. We identified 35 genes associated with blood lipids, including ten genes not previously shown to have gene-based signals. Our results support the hypothesis that genes closest to a GWAS index SNP are enriched for evidence of association.

## Supplemental data

Supplemental data includes in 8 figures, 49 tables, Study Descriptions, and Banner Authors.

## Declaration of Interests

The authors declare no competing interests for the present work. PN reports investigator-initiated grants from Amgen, Apple, and Boston Scientific, is a scientific advisor to Apple, Blackstone Life Sciences, and Novartis, and spousal employment at Vertex, all unrelated to the present work. A.V.K. has served as a scientific advisor to Sanofi, Medicines Company, Maze Pharmaceuticals, Navitor Pharmaceuticals, Verve Therapeutics, Amgen, and Color; received speaking fees from Illumina, MedGenome, Amgen, and the Novartis Institute for Biomedical Research; received sponsored research agreements from the Novartis Institute for Biomedical Research and IBM Research, and reports a patent related to a genetic risk predictor (20190017119). CJW spouse employed at Regeneron. Dr. Emery is currently an employee of Celgene/Bristol Myers Squibb. Celgene/Bristol Myers Squibb had no role in the funding, design, conduct, and interpretation of this study. MEM receives funding from Regeneron unrelated to this work. EEK has received speaker honoraria from Illumina, Inc and Regeneron Pharmceuticals. BMP serves on the Steering Committee of the Yale Open Data Access Project funded by Johnson & Johnson. LAC has consulted with the Dyslipidemia Foundation on lipid projects in the Framingham Heart Study. PTE is supported by a grant from Bayer AG to the Broad Institute focused on the genetics and therapeutics of cardiovascular disease. PTE has consulted for Bayer AG, Novartis, MyoKardia and Quest Diagnostics. SAL receives sponsored research support from Bristol Myers Squibb / Pfizer, Bayer AG, Boehringer Ingelheim, Fitbit, and IBM, and has consulted for Bristol Myers Squibb / Pfizer, Bayer AG, and Blackstone Life Sciences. The views expressed in this article are those of the author(s) and not necessarily those of the NHS, the NIHR, or the Department of Health. MMcC has served on advisory panels for Pfizer, NovoNordisk and Zoe Global, has received honoraria from Merck, Pfizer, Novo Nordisk and Eli Lilly, and research funding from Abbvie, Astra Zeneca, Boehringer Ingelheim, Eli Lilly, Janssen, Merck, NovoNordisk, Pfizer, Roche, Sanofi Aventis, Servier, and Takeda. As of June 2019, MMcC is an employee of Genentech, and a holder of Roche stock. MEJ holds shares in Novo Nordisk A/S. H.M.K. is an employee of Regeneron Pharmaceuticals; he owns stock and stock options for Regeneron Pharmaceuticals. MEJ has received research grants form Astra Zeneca, Boehringer Ingelheim, Amgen, Sanofi. SK is founder of Verve Therapeutics.

## Acknowledgements

This work was supported by a grant from the Swedish Research Council (2016-06830) to GH. This work was supported by grants from the National Heart, Lung, and Blood Institute (NHLBI): R01HL142711 and R01HL127564 (P.N. and G.M.P.), R03HL141439 (G.M.P.), R01HL148050 and R01HL148565 (PN). P.N. is also supported by a Hassenfeld Scholar Award from the Massachusetts General Hospital, Foundation Leducq (TNE-18CVD04), and additional grants from the National Heart, Lung, and Blood Institute (R01HL148565 and R01HL148050). PSdV is supported by American Heart Association grant number 18CDA34110116. BEC and JL are supported by R35HL135818, HL113338, and HL46380. BEC is also supported by K01HL135405. S.A.L. is supported by NIH grant 1R01HL139731 and American Heart Association 18SFRN34250007. AVK is supported by K08HG010155. JF is supported by R01DK125490. DJL is supported by R01HG008983 and R01GM126479. CJW is supported by R35HL135824, R01HL142023, and R01HL127564. MJB is supported British Heart Foundation CS/14/2/30841. OM is supported by European Research Council ERC-AdG-2019-885003, NNF17OC0026936, and VR 2018-02760. MOM is supported by the European Research Council ERC-CoG-2014-649021 and NNF18OC0034386. RM is supported by Canadian Institutes Health Research FDN 154308. SR is supported by R35HL135818, HL113338, HL46380, and K01HL135405. NDP is supported by R01 HL92301, R01 HL67348, R01 NS058700, R01 AR48797, R01 DK071891, R01 AG058921, the General Clinical Research Center of the Wake Forest University School of Medicine (M01 RR07122, F32 HL085989), the American Diabetes Association, and a pilot grant from the Claude Pepper Older Americans Independence Center of Wake Forest University Health Sciences (P60 AG10484). BIF is supported by R01 NS058700 and R01 DK071891. QQ is supported by R01-DK119268, R01-HL060712, and R01-HL140976. PTE is supported by the National Institutes of Health (1RO1HL092577, R01HL128914, K24HL105780), the American Heart Association (18SFRN34110082), and by the Foundation Leducq (14CVD01). NLHC is supported by HHSN268201500001. JCF is support by NIDDK U01 DK105554 and NIDDK K24 DK110550. MIM is supported by NIDDK U01-DK105535 and Wellcome 090532, 098381, 106130, 203141, 212259. NPB is supported by NIDDK U01-DK105554. NG is supported by Novo Nordisk Foundation Center for Basic Metabolic Research is an independent Research Center, based at the University of Copenhagen, Denmark and partially funded by an unconditional donation from the Novo Nordisk Foundation (www.cbmr.ku.dk) (Grant number NNF18CC0034900). TLT and HMM are supported by CONACYT, 312688. KLM is supported by US National Institutes of Health DK072193 and DK093757. CLH is supported by DK085591. EST is supported by the National Medical Research Council of Singapore Clinician Scientist Award. NB is supported by The American Federation for Aging Research, the Einstein Glenn Center and the NIA (PO1AG027734, R01AG 046949, 1R01AG042188 and P30AG038072). RCWM is supported by the Hong Kong Research Grants Council Theme-based Research Scheme (T12-402/13N) and Research Impact Fund (R4012-18). The KARE cohort was supported by grants from Korea Centers for Disease Control and Prevention (4845–301, 4851–302, 4851–307) and intramural grants from the Korea National Institute of Health (2019-NG-053-01). Collection of the San Antonio Family Study data were supported in part by National Institutes of Health (NIH) grants P01 HL045522, R01 MH078143, R01 MH078111 and R01 MH083824; and whole genome sequencing of SAFS subjects was supported by U01 DK085524 and R01 HL113323. We are very grateful to the participants of the San Antonio Family Study for their continued involvement in our research programs. DGI (Principal investigators Leif Groop and Tiinamaija Tuomi) and the Botnia Study is supported by the Sigrid Juselius Foundation, The Folkhalsan Research Foundation, Nordic Center of Excellence in Disease Genetics, EU (EXGENESIS), Finnish Diabetes Research Foundation, Foundation for Life and Health in Finland, Finnish Medical Society, Helsinki University Central Hospital Research Foundation, Perklén Foundation, Ollqvist Foundation, Narpes Health Care Foundation as well as the Municipal Heath Care Center and Hospital in Jakobstad and Health Care Centers in Vasa, Narpes and Korsholm. The work in Malmö, Sweden, was also funded by a Linné grant from the Swedish Research Council (349-2006-237). The contribution of the Botnia and Skara research teams is gratefully acknowledged. The TwinsUK study was funded by the Wellcome Trust and European Community’s Seventh Framework Programme (FP7/2007-2013). The TwinsUK study also receives support from the National Institute for Health Research (NIHR)-funded BioResource, Clinical Research Facility and Biomedical Research Centre based at Guy’s and St Thomas’ NHS Foundation Trust in partnership with King’s College London. The KORA study was initiated and financed by the Helmholtz Zentrum München – German Research Center for Environmental Health, which is funded by the German Federal Ministry of Education and Research (BMBF) and by the State of Bavaria. Furthermore, KORA research was supported within the Munich Center of Health Sciences (MC-Health), Ludwig-Maximilians-Universität, as part of LMUinnovativ. Funded by the Bavarian State Ministry of Health and Care through the research project DigiMed Bayern (www.digimed-bayern.de).

Molecular data for the Trans-Omics in Precision Medicine (TOPMed) program was supported by the National Heart, Lung and Blood Institute (NHLBI). See the TOPMed Omics Support information in the **Supplementary Text** for study specific omics support information. Core support including centralized genomic read mapping and genotype calling, along with variant quality metrics and filtering were provided by the TOPMed Informatics Research Center (3R01HL-117626-02S1; contract HHSN268201800002I). Core support including phenotype harmonization, data management, sample-identity QC, and general program coordination were provided by the TOPMed Data Coordinating Center (R01HL-120393; U01HL-120393; contract HHSN268201800001I). We gratefully acknowledge the studies and participants who provided biological samples and data for TOPMed. The views expressed in this manuscript are those of the authors and do not necessarily represent the views of the National Heart, Lung, and Blood Institute; the National Institutes of Health; or the U.S. Department of Health and Human Services.

## Data and code availability

Controlled access of the individual-level data are available through dbGAP (please refer to the Supplementary Information), and the individual-level UK Biobank data are available upon application to the UK Biobank

