## Supplemental Text and Figures for "Rare coding variants in 35 genes associate with circulating lipid levels – a multi-ancestry analysis of 170,000 exomes"

**Hindy et al.**

**Supplementary Information**

**Figure S1. Comparison of effect sizes and p-value in UK Biobank including and excluding individuals on statin treatment.**

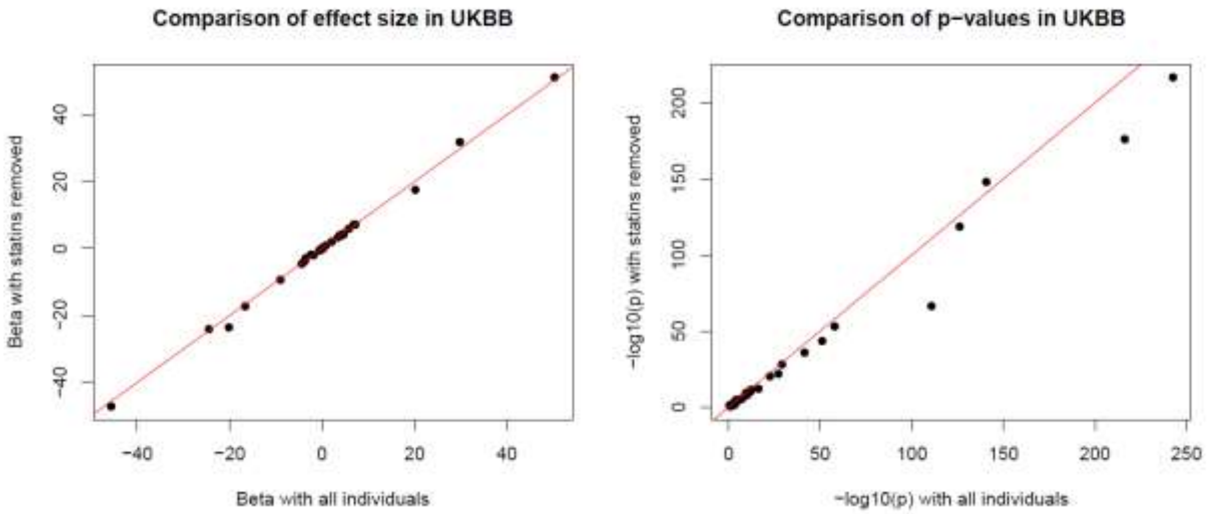

**Figure S2. Descriptive variant characteristics by type, ancestry, and minor allele count.**

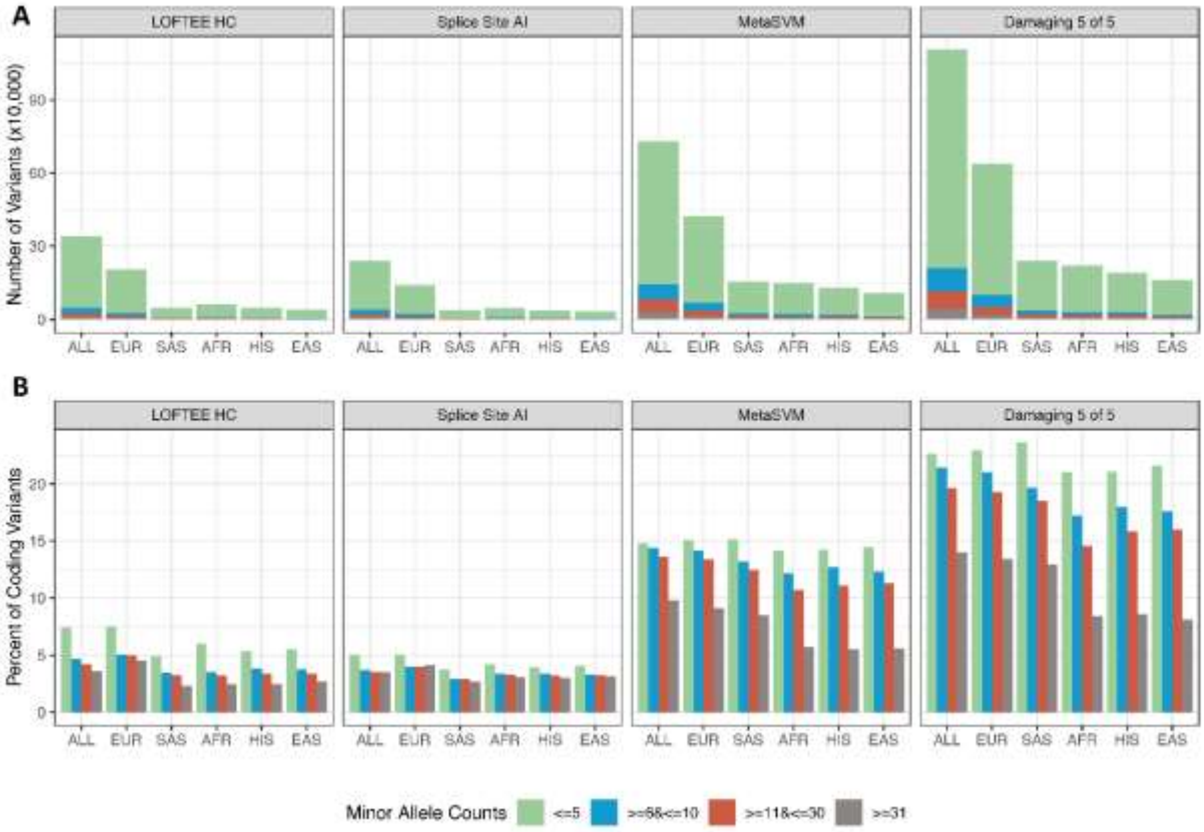


**Figure S2: A)** Our study included 15,599,513 genetic variants. Variants were annotated as high confidence loss-of-function by LOFTEE (n=340,214), splice site altering variants using a deep neural network prediction (SPLICE AI) (n=238,646), damaging missense variants according to the MetaSVM algorithm (n=729,098) and damaging missing in 5 out of 5 prediction algorithms (n=1,106,309). Most of the variants had a minor allele count of less than 5 in all (n=1,171,5189) and within each of the four different annotations. **B)** The proportion of specific annotations out of the total number of variants that were annotated as coding (n=5,085,712). Each of the four annotations demonstrated the highest enrichment among the variants with the lowest frequency. ALL=multi-ancestry, AFR=African ancestry, EAS=East Asian ancestry, EUR=European ancestry, HIS=Hispanic ancestry, SAS=South Asian ancestry.

**Figure S3. Overlap among different ancestries for all variants contributing to significant gene-based associations with HDL-C, TG and LDL-C**

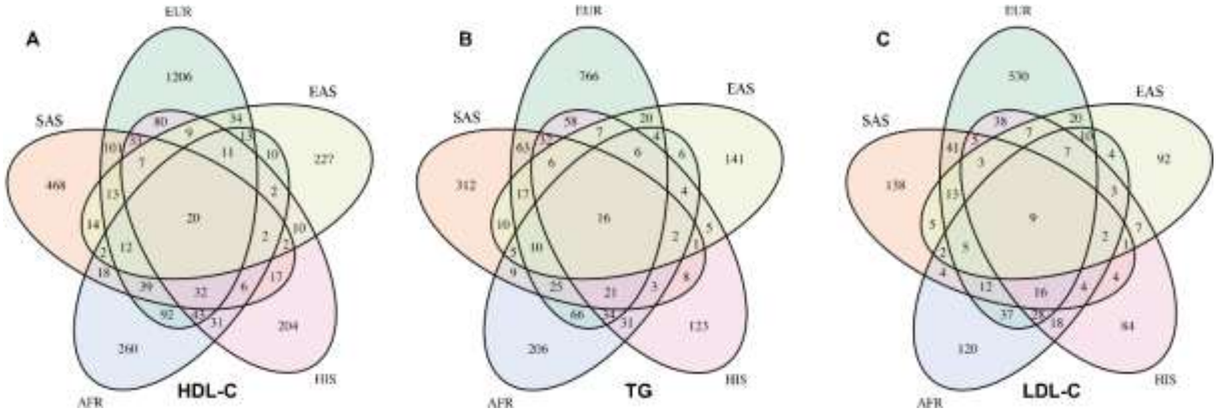


**Figure S3:** Venn Diagram for the overlap of all variants included in the significant gene-based association analysis among different ancestries. **A)** A total of 15 genes showed significant gene-based associations in multi-ancestry analyses with HDL cholesterol (HDL-C). Ancestry-specific single-variant contributions included a total of 1745 from European-, 786 from South Asian-, 593 African-, 509 Hispanic- and 388 East Asian ancestries. **B)** A total of 17 genes showed significant gene-based associations in multi-ancestry analyses with triglycerides (TG). Ancestry-specific single-variant contributions included a total of 1151 from European-, 540 from South Asian-, 448 African-, 357 Hispanic- and 260 East Asian ancestries. **C)** A total of 8 genes showed significant gene-based associations in multi-ancestry analyses with triglycerides (TG). Ancestry-specific single-variant contributions included a total of 1086 from European-, 396 from South Asian-, 371 African-, 292 Hispanic- and 280 East Asian ancestries.

**Figure S4. Overlap among different ancestries for variants contributing to significant gene-based associations with HDL-C, TG and LDL-C with P value less than 0.05**

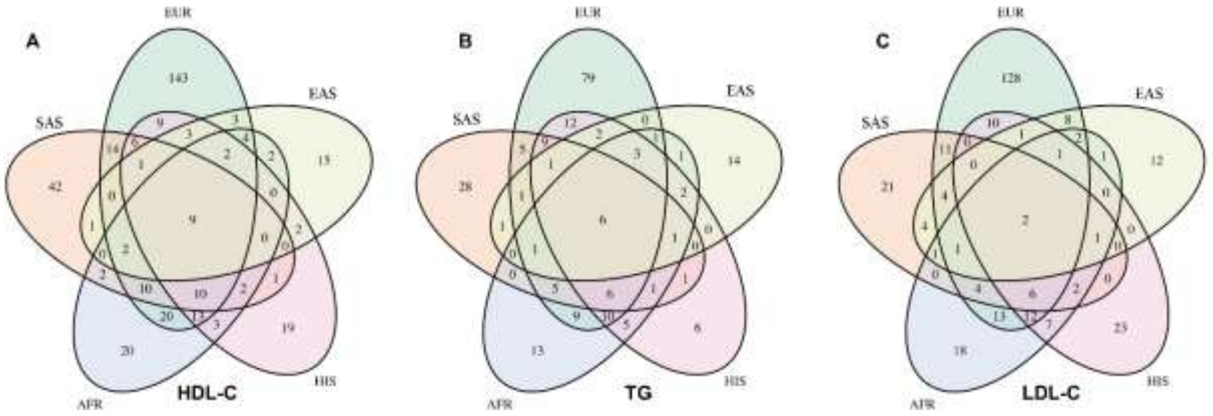


**Figure S4:** Venn Diagram for the overlap of variants with P<0.05 included in the significant gene-based association analysis among different ancestries. **A)** A total of 15 genes showed significant gene-based associations in multi-ancestry analyses with HDL cholesterol (HDL-C). Ancestry-specific single-variant contributions included a total of 249 from European-, 100 from South Asian-, 99 African-, 80 Hispanic- and 44 East Asian ancestries. **B)** A total of 17 genes showed significant gene-based associations in multi-ancestry analyses with triglycerides (TG). Ancestry-specific single-variant contributions included a total of 150 from European-, 66 from South Asian-, 64 African-, 65 Hispanic- and 34 East Asian ancestries. **C)** A total of 8 genes showed significant gene-based associations in multi-ancestry analyses with triglycerides (TG). Ancestry-specific single-variant contributions included a total of 331 from European-, 116 from South Asian-, 114 African-, 99 Hispanic- and 73 East Asian ancestries.

**Figure S5. Overlap among different ancestries for the top variant contributing to significant gene-based associations with HDL-C, TG and LDL-C**

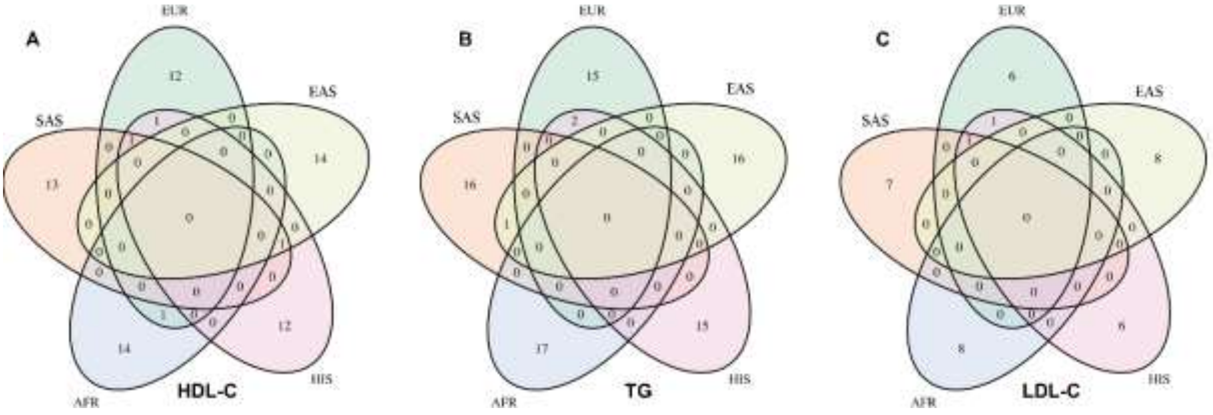


**Figure S5:** Venn Diagram for the overlap of the top variant included in each of the significant gene-based association analysis among different ancestries. **A)** A total of 15 genes showed significant gene-based associations in multi-ancestry analyses with HDL cholesterol (HDL-C). **B)** A total of 17 genes showed significant gene-based associations in multi-ancestry analyses with triglycerides (TG). **C)** A total of 8 genes showed significant gene-based associations in multi-ancestry analyses with triglycerides (TG).

**Figure S6. Cumulative loss-of-function minor allele count and effect size on LDL cholesterol by ancestry**

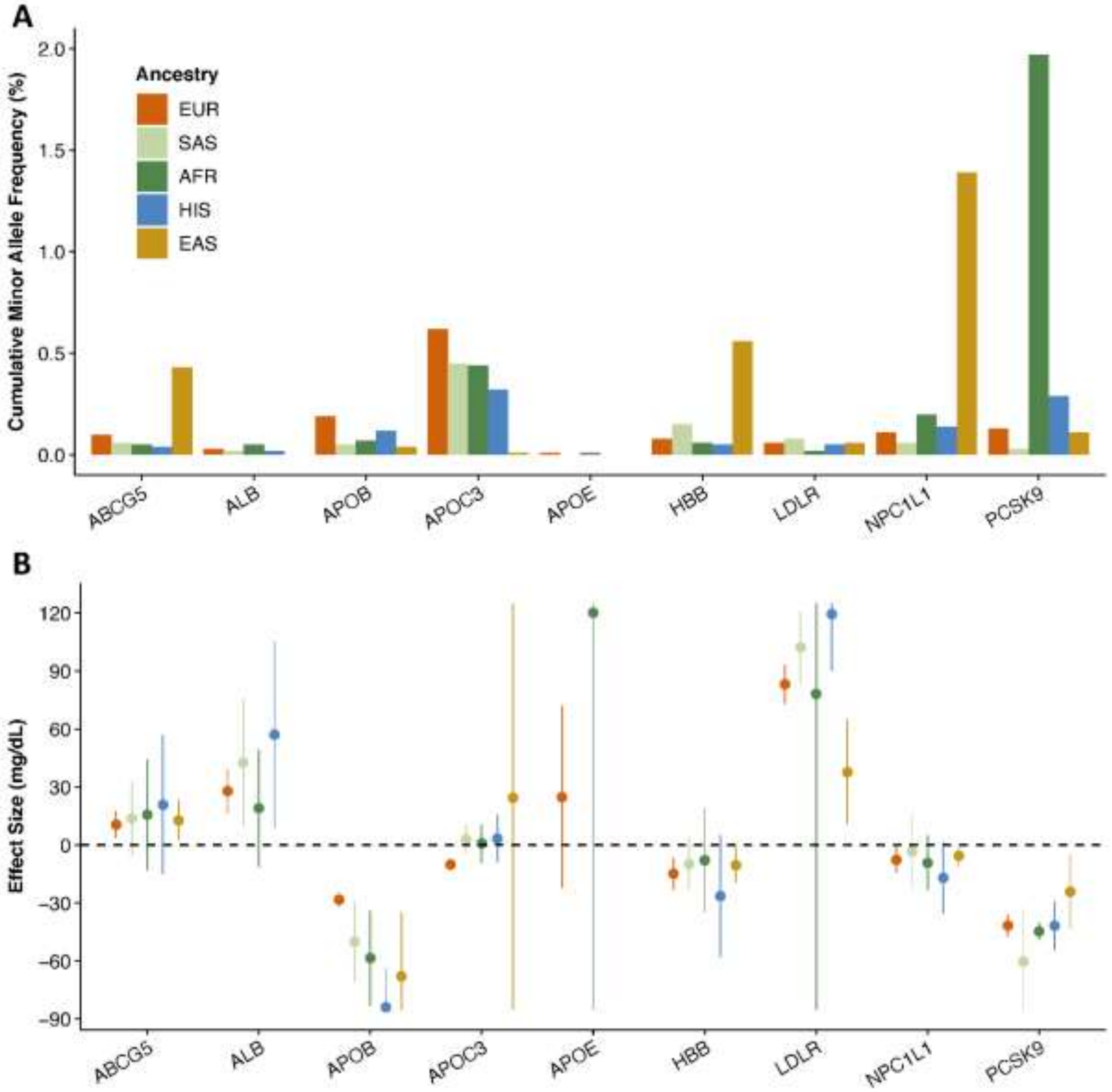


**Figure S6: A)** Cumulative minor allele frequencies and **B)** burden test effect sizes on LDL cholesterol levels for exome-wide significant genes ($P < 4.3\times10^{-7}$) within each of the five major ancestries using variants from the high confidence loss-of-function grouping (LOFTEE). AFR=African ancestry, EAS=East Asian ancestry, EUR=European ancestry, HIS=Hispanic ancestry, SAS=South Asian ancestry.

**Figure S7. Cumulative loss-of-function minor allele count and effect size on triglycerides by ancestry**

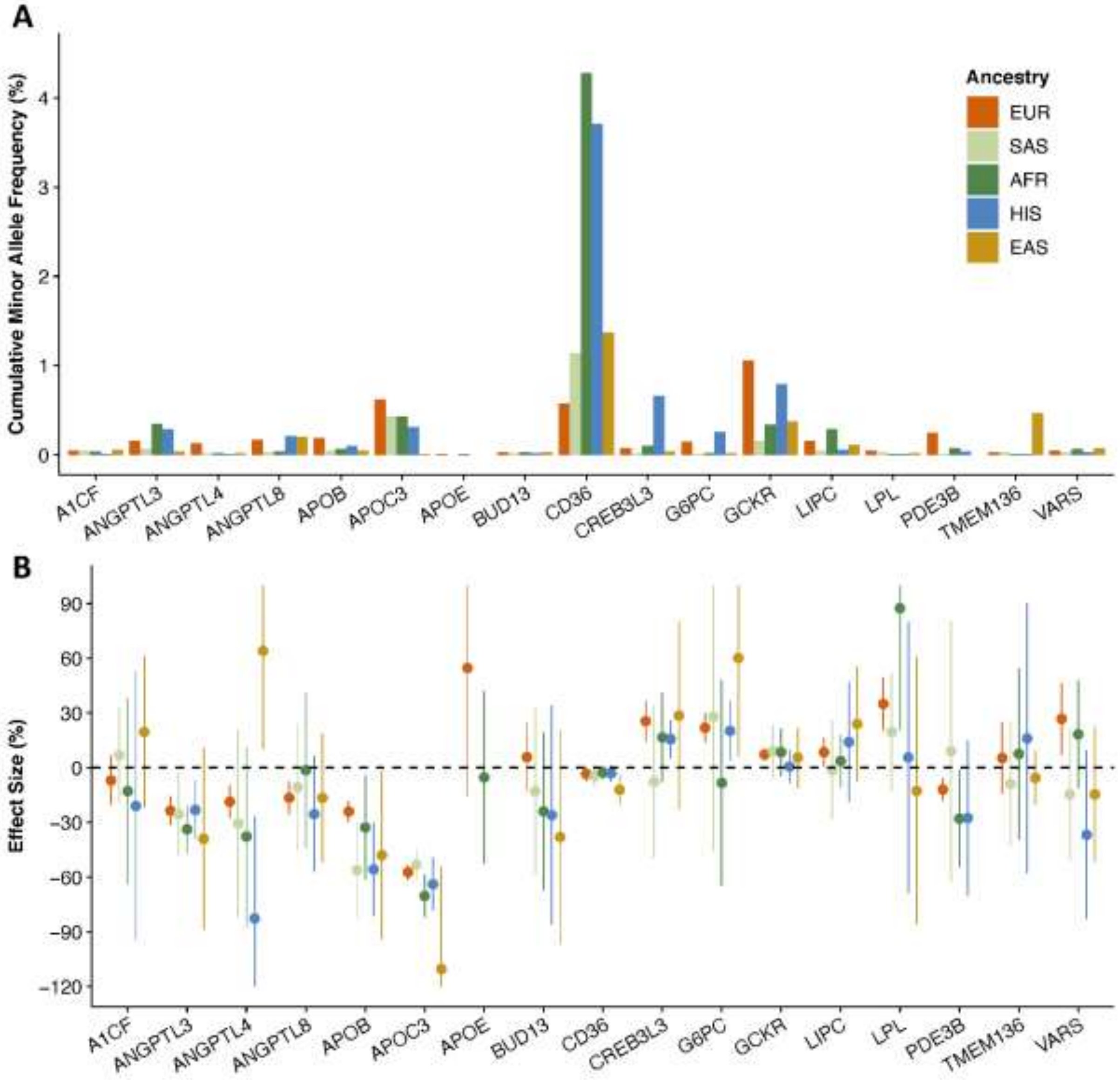


**Figure S7: A)** Cumulative minor allele frequencies and **B)** burden test effect sizes on triglyceride levels for exome-wide significant genes ($P < 4.3\times10^{-7}$) within each of the five major ancestries using variants from the high confidence loss-of-function grouping (LOFTEE). AFR=African ancestry, EAS=East Asian ancestry, EUR=European ancestry, HIS=Hispanic ancestry, SAS=South Asian ancestry.

**Figure S8. Cumulative loss-of-function minor allele count and effect size on HDL cholesterol by ancestry**

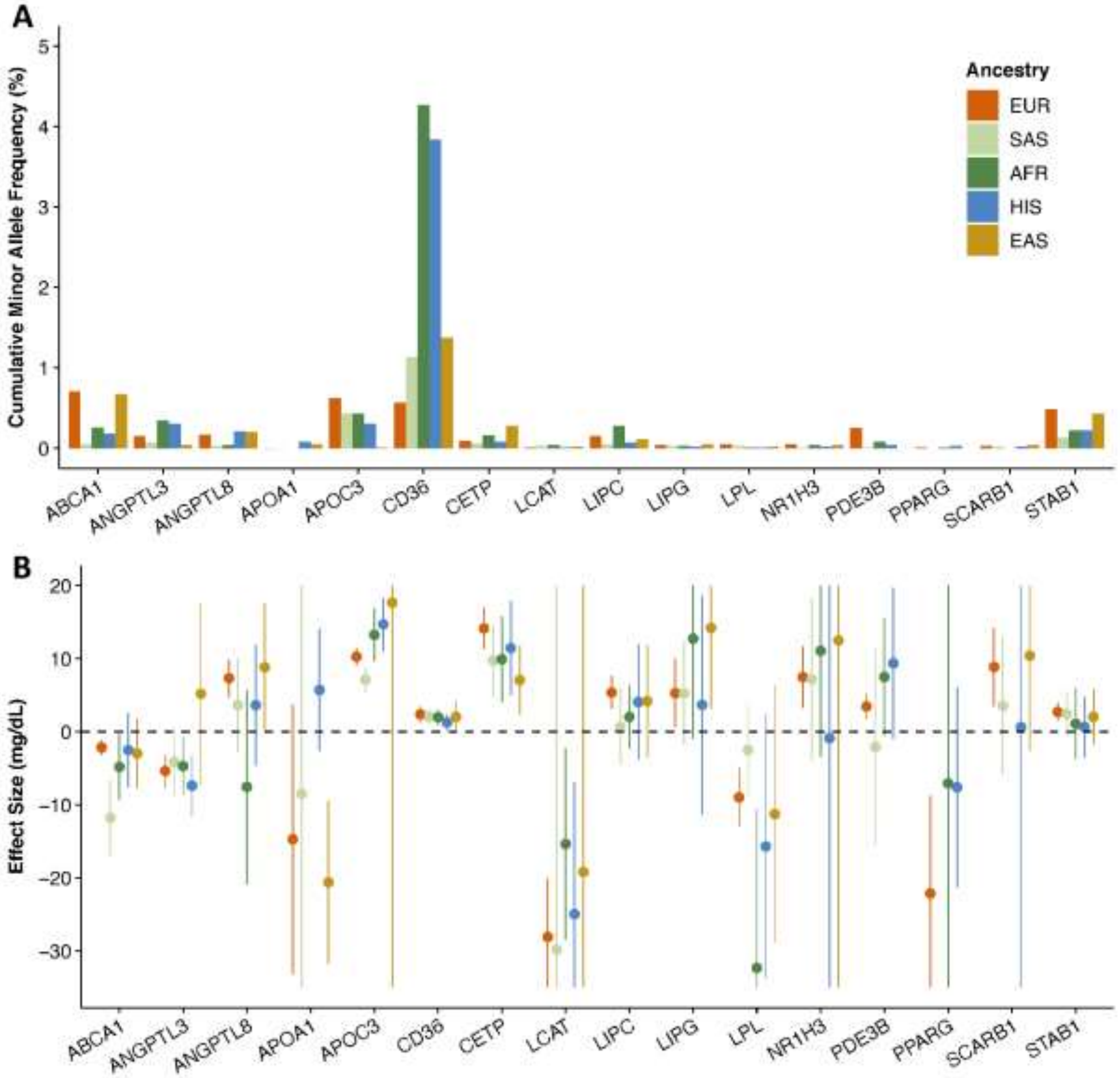


**Figure S8: A)** Cumulative minor allele frequencies and **B)** burden test effect sizes on HDL cholesterol levels for exome-wide significant genes ($P < 4.3 \times 10^{-7}$) within each of the five major ancestries using variants from the high confidence loss-of-function grouping (LOFTEE). AFR=African ancestry, EAS=East Asian ancestry, EUR=European ancestry, HIS=Hispanic ancestry, SAS=South Asian ancestry.

### Study Participant Descriptions

#### Myocardial Infarction Genetics Consortium (MIGen) study participants

MIGen studies included the Atherosclerosis Risk in Communities study (ARIC), Italian Atherosclerosis Thrombosis and Vascular Biology (ATVB) study,[1] Bangladesh Risk of Acute Vascular Events study (BRAVE),[2] the Exome Sequencing Project Early-Onset Myocardial Infarction (ESP-EOMI) study,[3] a nested case-control cohort of the Jackson Heart Study (JHS),[4] the South German Myocardial Infarction study,[5] the Ottawa Heart Study (OHS),[6] the Precocious Coronary Artery Disease Study (PROCARDIS),[7] the Pakistan Risk of Myocardial Infarction Study (PROMIS),[8] the Registre Gironi del COR (Gerona Heart Registry or REGICOR) study,[9] the Leicester Myocardial Infarction study,[10] and the North German Myocardial Infarction study[11] (**Supplemental Table 37**). Clinical data were assessed in each study.

All participants in the study provided written informed consent for genetic studies. The institutional review boards at the Broad Institute and each participating institution approved the study protocol.

In order to minimize the possibility of unintentionally sharing information that can be used to re-identify private information, a subset of the data generated for this study are available at dbGaP and can be accessed at through dbGaP Study Accessions: phs000090.v1.p1 (ARIC), phs000814.v1.p1 (ATVB), phs001398.v1.p1 (BRAVE), phs000279.v2.p1 (EOMI), phs001098.v1.p1 (JHS), phs001000.v1.p1 (Leicester), phs000990.v1.p1 (NorthGermanMI), phs000916.v1.p1 (SouthGermanMI), phs000806.v1.p1 (OHS), phs000883.v1.p1 (PROCARDIS), phs000917.v1.p1 (PROMIS), phs000902.v1.p1(Regicor).

#### TOPMed program study participants

<u>Atherosclerosis Risk in Communities study (ARIC, 2868)</u>

*TOPMed dbGaP accession#: phs001211, Parent dbGaP accession#: phs000280*

ARIC is a large population-based prospective longitudinal cohort study (began 1987) from four U.S. communities: Forsyth County, NC; Jackson, MS; the northwest suburbs of Minneapolis, MN; and Washington County, MD. ARIC was designed to investigate the etiology and natural history of atherosclerosis, its consequences, and related medical care by race, gender, location, and time as previously described.[12] A total of 15,792 participants (55% female and 27% African American) aged 45-64 years were recruited between 1987 and 1989 and received extensive examination, including medical, social and demographic data. The baseline visit was conducted between 1987 and1989, the second visit in 1990-1992, the third visit in 1993-1995, the fourth visit in 1996-1998, the fifth visit in 2011-2013, the sixth visit in 2016-2017, and the seventh visit in 2018-2019. Follow-up is also conducted semi-annually since 2012 (annually prior to that) by telephone to maintain contact with participants and to assess the health status of the cohort.

The Atherosclerosis Risk in Communities study has been funded in whole or in part with Federal funds from the National Heart, Lung, and Blood Institute, National Institutes of Health, Department of Health and Human Services (contract numbers HHSN268201700001I, HHSN268201700002I, HHSN268201700003I, HHSN268201700004I and HHSN268201700005I). The authors thank the staff and participants of the ARIC study for their important contributions.

Whole genome sequencing (WGS) for the Trans-Omics in Precision Medicine (TOPMed) program was supported by the National Heart, Lung and Blood Institute (NHLBI). WGS for “NHLBI TOPMed: Atherosclerosis Risk in Communities (ARIC)” (phs001211) was performed at the Baylor College of Medicine Human Genome Sequencing Center (HHSN268201500015C and 3U54HG003273-12S2) and the Broad Institute for MIT and Harvard (3R01HL092577-06S1). Centralized read mapping and genotype calling, along with variant quality metrics and filtering were provided by the TOPMed Informatics Research Center (3R01HL-117626-02S1). Phenotype harmonization, data management, sample-identity QC, and general study coordination, were provided by the TOPMed Data Coordinating Center (3R01HL-120393-02S1). We gratefully acknowledge the studies and participants who provided biological samples and data for TOPMed.

The Genome Sequencing Program (GSP) was funded by the National Human Genome Research Institute (NHGRI), the National Heart, Lung, and Blood Institute (NHLBI), and the National Eye Institute (NEI). The GSP Coordinating Center (U24 HG008956) contributed to cross-program scientific initiatives and provided logistical and general study coordination. The Centers for Common Disease Genomics (CCDG) program was supported by NHGRI and NHLBI, and whole genome sequencing was performed at the Baylor College of Medicine Human Genome Sequencing Center (UM1 HG008898 and R01HL059367).

Old Order Amish (Amish, 1,083)

*TOPMed dbGaP accession#: phs000956, Parent dbGaP accession#: phs000391*

The Amish Complex Disease Research Program includes a set of large community-based studies focused largely on cardiometabolic health carried out in the Old Order Amish (OOA) community of Lancaster, Pennsylvania.[13] The OOA population of Lancaster County, PA immigrated to the Colonies from Western Europe in the early 1700's. There are now over 30,000 OOA individuals in the Lancaster area, nearly all of whom can trace their ancestry back 12-14 generations to approximately 700 founders. Investigators at the University of Maryland School of Medicine have been studying the genetic determinants of cardiometabolic health in this population since 1993. To date, over 7,000 Amish adults have participated in one or more of our studies.

The Amish studies upon which these data are based were supported by NIH grants R01 AG18728, U01 HL072515, R01 HL088119, R01 HL121007, U01 HL137181, and P30 DK072488, American Heart Association grant AHA 17GRNT33661168 WGS for “NHLBI TOPMed: Genetics of Cardiometabolic Health in the Amish” (phs000956) was performed at the Broad Institute of MIT and Harvard (3R01HL121007-01S1).

Mt Sinai BioMe Biobank (BioMe, 3257)

*TOPMed dbGaP accession#: phs001644, Parent dbGaP accession#: phs000925*

The Mount Sinai Institute for Personalized Medicine BioMe Biobank is a consented, EMR-linked medical care setting biorepository of the Mount Sinai Medical Center drawing from a population of over 70,000 inpatients and 800,000 outpatient visits annually.[14] The Mount Sinai Medical Center services diverse local communities of upper Manhattan, including Central Harlem (86% African American), East Harlem

(88% Hispanic Latino), and Upper East Side (88% European ancestry/white) with broad health disparities. Biobank operations are fully integrated in clinical care processes, including direct recruitment from clinical sites waiting areas and phlebotomy stations by dedicated Biobank recruiters independent of clinical care providers, prior to or following a clinician standard of care visit. Recruitment currently occurs at a broad spectrum of over 30 clinical care sites.

The Mount Sinai BioMe Biobank has been supported by The Andrea and Charles Bronfman Philanthropies and in part by Federal funds from the NHLBI and NHGRI (U01HG00638001; U01HG007417; X01HL134588). WGS for “NHLBI TOPMed: Mount Sinai BioMe Biobank” (phs001644) was performed at the Baylor College of Medicine Human Genome Sequencing Center (HHSN268201600033I). We thank all participants in the Mount Sinai Biobank. We also thank all our recruiters who have assisted and continue to assist in data collection and management and are grateful for the computational resources and staff expertise provided by Scientific Computing at the Icahn School of Medicine at Mount Sinai.

<u>Coronary Artery Risk Development in Young Adults (CARDIA, 2724)</u>

*TOPMed dbGaP accession#: phs001612, Parent dbGaP accession#: phs000285*

The Coronary Artery Risk Development in Young Adults Study (CARDIA) is a study examining the etiology and natural history of cardiovascular disease beginning in young adulthood.[15] In 1985-1986, a cohort of 5115 healthy black and white men and women aged 18-30 years were selected to have approximately the same number of people in subgroups of age (18-24 and 25-30), sex, race, and education (high school or less and more than high school) within each of four US Field Centers. These same participants were asked to participate in follow-up examinations during 1987-1988 (Year 2), 1990-1991 (Year 5), 1992-1993 (Year 7), 1995-1996 (Year 10), 2000-2001 (Year 15), 2005-2006 (Year 20), 2010-2011 (Year 25); and 2015-2016 (Year 30). A majority of the group has been examined at each of the follow-up examinations (91%, 86%, 81%, 79%, 74%, 72%, 72%, and 71%, respectively). In addition to the follow-up examinations, participants are contacted regularly for the ascertainment of information on out-patient procedures and hospitalizations experienced between contacts.

The Coronary Artery Risk Development in Young Adults Study (CARDIA) is conducted and supported by the National Heart, Lung, and Blood Institute (NHLBI) in collaboration with the University of Alabama at Birmingham (HHSN268201800005I & HHSN268201800007I), Northwestern University (HHSN268201800003I), University of Minnesota (HHSN268201800006I), and Kaiser Foundation Research Institute (HHSN268201800004I). CARDIA was also partially supported by the Intramural Research Program of the National Institute on Aging (NIA) and an intra-agency agreement between NIA and NHLBI (AG0005). WGS for “NHLBI TOPMed: Coronary Artery Risk Development in Young Adults” (phs001612) was performed at the Baylor College of Medicine Human Genome Sequencing Center (HHSN268201600033I).

<u>Cleveland Family Study (CFS, 532)</u>

*TOPMed dbGaP accession#: phs000954, Parent dbGaP accession#: phs000284*

The Cleveland Family Study (CFS) is a family-based study of sleep apnea, comprising of 2,284 individuals (46% African American) from 361 families studied up to 4 occasions over 16 years, 1990-2006.[16-19] Index probands (n=275) were recruited from 3 area hospital sleep labs if they had a confirmed diagnosis of sleep apnea and at least 2 first-degree relatives available to be studied. In the first 5 years of the study, neighborhood control probands (n=87) with at least 2 living relatives available for study were selected at random from a list provided by the index family and also studied. All available first-degree relatives and spouses of the case and control probands also were recruited. Second-degree relatives, including half-sibs, aunts, uncles and grandparents, were also included if they lived near the first-degree relatives (cases or controls), or if the family had been found to have two or more relatives with sleep apnea. Blood was sampled and DNA isolated for participants seen in the last two exam cycles (n=1,447).

CFS is supported by grants from the NHLBI (HL046389, HL113338, and 1R35HL135818). WGS for “NHLBI TOPMed: Cleveland Family Study - WGS Collaboration” (phs000954) was performed at the University of Washington Northwest Genomics Center (3R01HL098433-05S1 and HHSN268201600032I).

Cardiovascular Health Study (CHS, 2070)

*TOPMed dbGaP accession#: phs001368, Parent dbGaP accession#: phs000287*

The Cardiovascular Health Study (CHS) originated in 1988 and is a study of risk factors for development and progression of coronary heart disease and stroke in people aged 65 years and older.[20-22] The 5,888 study participants were recruited from four U.S. communities and have undergone extensive clinic examinations for evaluation of markers of subclinical cardiovascular disease. The original cohort totaled 5,201 participants. A new cohort was recruited in 1992. The 687 participants in the new cohort are predominately African-American and were recruited at three of the four field centers. Starting in 1989, and continuing through 1999, participants underwent annual extensive clinical examinations. Measurements included traditional risk factors such as blood pressure and lipids as well as measures of subclinical disease, including echocardiography of the heart, carotid ultrasound, and cranial magnetic-resonance imaging (MRI). At six-month intervals between clinic visits, and once clinic visits ended, participants were contacted by phone to ascertain hospitalizations and health status. The main outcomes are coronary heart disease (CHD), angina, heart failure (HF), stroke, transient ischemic attack (TIA), claudication, and mortality. Participants continue to be contacted by phone every 6 months.

This CHS research was supported by NHLBI contracts HHSN268201200036C, HHSN268200800007C, HHSN268201800001C, N01HC55222, N01HC85079, N01HC85080, N01HC85081, N01HC85082, N01HC85083, N01HC85086; and NHLBI grants U01HL080295, R01HL087652, R01HL105756, R01HL103612, R01HL120393, R01HL130114, and R01 HL059367, with additional contribution from the National Institute of Neurological Disorders and Stroke (NINDS). Additional support was provided through R01AG023629 from the National Institute on Aging (NIA). A full list of principal CHS investigators and institutions can be found at CHS-NHLBI.org. WGS for “NHLBI TOPMed: Cardiovascular Health Study” (phs001368) was performed at the Baylor College of Medicine Human Genome Sequencing Center (3U54HG003273-12S2,

HHSN268201500015C, and HHSN268201600033I). The content is solely the responsibility of the authors and does not necessarily represent the official views of the National Institutes of Health.

Diabetes Heart Study (DHS, 345)

*TOPMed dbGaP accession#: phs001412, Parent dbGaP accession#: phs001012*

The Diabetes Heart Study (DHS) is a family-based study enriched for type 2 diabetes (T2D).[23] The cohort included 1443 European American and African American participants from 564 families with multiple cases of type 2 diabetes. The cohort was recruited between 1998 and 2006. Participants were extensively phenotyped for measures of subclinical CVD and other known CVD risk factors. Primary outcomes were quantified burden of vascular calcified plaque in the coronary artery, carotid artery, and abdominal aorta all determined from non-contrast computed tomography scans.

This work was supported by R01 HL92301, R01 HL67348, R01 NS058700, R01 AR48797, R01 DK071891, R01 AG058921, the General Clinical Research Center of the Wake Forest University School of Medicine (M01 RR07122, F32 HL085989), the American Diabetes Association, and a pilot grant from the Claude Pepper Older Americans Independence Center of Wake Forest University Health Sciences (P60 AG10484). WGS for “NHLBI TOPMed: Diabetes Heart Study” (phs001412) was performed at the Broad Institute of MIT and Harvard (HHSN268201500014C).

Framingham Heart Study (FHS, 3,961)

*TOPMed dbGaP accession#: phs000974, Parent dbGaP accession#: phs000007*

The Framingham Heart Study (FHS) is a prospective cohort study of 3 generations of subjects who have been followed up to 65 years to evaluate risk factors for cardiovascular disease.[24-27] Its large sample of ~15,000 men and women who have been extensively phenotyped with repeated examinations make it ideal for the study of genetic associations with cardiovascular disease risk factors and outcomes. DNA samples have been collected and immortalized since the mid-1990s and are available on ~8000 study participants in 1037 families. These samples have been used for collection of GWAS array data and exome chip data in nearly all with DNA samples, and for targeted sequencing, deep exome sequencing and light coverage whole genome sequencing in limited numbers. Additionally, mRNA and miRNA expression data, DNA methylation data, metabolomics and other 'omics data are available on a sizable portion of study participants. This project will focus on deep whole genome sequencing (mean 30X coverage) in ~4100 subjects and imputed to all with GWAS array data to more fully understand the genetic contributions to cardiovascular, lung, blood and sleep disorders.

FHS acknowledges the support of contracts NO1-HC-25195 and HHSN268201500001I from the National Heart, Lung and Blood Institute and grant supplement R01 HL092577-06S1 for this research. WGS for “NHLBI TOPMed: Whole Genome Sequencing and Related Phenotypes in the Framingham Heart Study” (phs000974) was performed at the Broad Institute of MIT and Harvard (HHSN268201500014C, 3R01HL092577-06S1, and 3U54HG003067-12S2). We also acknowledge the dedication of the FHS study participants without whom this research would not be possible.

Genetic Epidemiology Network of Arteriopathy (GENOA, 391)

*TOPMed dbGaP accession#: phs001345, Parent dbGaP accession#: phs001238*

The Genetic Epidemiology Network of Arteriopathy (GENOA) is one of four networks in the NHLBI Family-Blood Pressure Program (FBPP).28 GENOA's long-term objective is to elucidate the genetics of target organ complications of hypertension, including both atherosclerotic and arteriolosclerotic complications involving the heart, brain, kidneys, and peripheral arteries.29 The longitudinal GENOA Study recruited European-American and African-American sibships with at least 2 individuals with clinically diagnosed essential hypertension before age 60 years. All other members of the sibship were invited to participate regardless of their hypertension status. Participants were diagnosed with hypertension if they had either 1) a previous clinical diagnosis of hypertension by a physician with current anti-hypertensive treatment, or 2) an average systolic blood pressure ≥ 140 mm Hg or diastolic blood pressure ≥ 90 mm Hg based on the second and third readings at the time of their clinic visit. Only participants of the African-American Cohort were sequenced through TOPMed.

Support for GENOA was provided by the National Heart, Lung and Blood Institute (HL054457, HL054464, HL054481, and HL087660) of the National Institutes of Health. WGS for “NHLBI TOPMed: Genetic Epidemiology Network of Arteriopathy” (phs001345) was performed at the Broad Institute of MIT and Harvard (HHSN268201500014C) and the University of Washington Northwest Genomics Center (3R01HL055673-18S1).

Genetics of Lipid-Lowering Drugs and Diet Network (GOLDN, 594)

*TOPMed dbGaP accession#: phs001359, Parent dbGaP accession#: phs000741*

The Genetics of Lipid-Lowering Drugs and Diet Network (GOLDN) study was initiated to assess how genetic factors interact with environmental (diet and drug) interventions to influence blood levels of triglycerides and other atherogenic lipid species and inflammation markers (registered at clinicaltrials.gov, number NCT00083369).[30] The study recruited participants of European ancestry primarily from three-generational pedigrees from two NHLBI Family Heart Study (FHS) field centers (Minneapolis, MN and Salt Lake City, UT).[31] Only families with at least two siblings were recruited and only participants who did not take lipid-lowering agents (pharmaceuticals or nutraceuticals) for at least 4 weeks prior to the initial visit were included. The diet intervention followed the protocol of Patsch et al.[32] The whipping cream (83% fat) meal had 700 Calories/m2 body surface area (2.93 mJ/m2 body surface area): 3% of calories were derived from protein (instant nonfat dry milk) and 14% from carbohydrate (sugar). The ratio of polyunsaturated to saturated fat was 0.06 and the cholesterol content of the average meal was 240 mg. The mixture was blended with ice and flavorings. Blood samples were drawn immediately before (fasting) and at 3.5 and 6 hours after consuming the high-fat meal. The diet intervention was administered at baseline as well as after a 3-week treatment with 160 mg micronized fenofibrate. Participants were given the option to complete one or both (diet and drug) interventions. Of all participants, 1079 had phenotypic data and provided appropriate consent, and underwent whole genome sequencing through the TOPMed program.

GOLDN biospecimens, baseline phenotype data, and intervention phenotype data were collected with funding from National Heart, Lung and Blood Institute (NHLBI) grant U01 HL072524. WGS for "NHLBI TOPMed: Genetics of Lipid Lowering Drugs and Diet Network" (phs001359) was performed at the University of Washington Northwest Genomics Center (3R01HL104135-04S1 and R01 HL104135).

Genetic Epidemiology Network of Salt Sensitivity (GenSalt, 1,749)

*TOPMed dbGaP accession#: phs001217, Parent dbGaP accession#: phs000784*

The Genetic Epidemiology Network of Salt-Sensitivity (GenSalt) study, using a family feeding-study design, aims to identify genes which interact with dietary sodium and potassium intake to influence blood pressure in Han Chinese participants from rural north China.[33] The dietary intervention included a 7-day low-sodium feeding (51.3 mmol/day), a 7-day high-sodium feeding (307.8 mmol/day) and a 7-day high-sodium feeding with an oral potassium supplementation (60 mmol/day). Microsatellite markers for genome-wide linkage scan and single nucleotide polymorphism (SNP) markers in candidate genes will be genotyped. Overall, 3153 participants from 658 families were recruited for GenSalt. Whole genome sequencing has been conducted for 1,860 participants as a part of TOPMed.

GenSalt was supported by research grants (U01HL072507, R01HL087263, and R01HL090682) from the National Heart, Lung and Blood Institute, National Institutes of Health, Bethesda, MD. WGS for "NHLBI TOPMed: Genetic Epidemiology Network of Salt Sensitivity" (phs001217) was performed at the Baylor College of Medicine Human Genome Sequencing Center (HHSN268201500015C).

Genetic Studies of Atherosclerosis Risk (GeneSTAR, 1,749)

*TOPMed dbGaP accession#: phs001218, Parent dbGaP accession#: phs000375*

GeneSTAR began in 1982 as the Johns Hopkins Sibling and Family Heart Study, a prospective longitudinal family-based study conducted originally in healthy adult siblings of people with documented early onset coronary disease under 60 years of age.[34,35] Commencing in 2003, the siblings, their offspring, and the coparent of the offspring participated in a 2 week trial of aspirin 81 mg/day with pre and post ex vivo platelet function assessed using multiple agonists in whole blood and platelet rich plasma. Extensive additional cardiovascular testing and risk assessment was done at baseline and serially. Follow-up was carried out to determine incident cardiovascular disease, stroke, peripheral arterial disease, diabetes, cancer, and related comorbidities, from 5 to 30 years after study entry. The goal of several additional phenotyping and interventional substudies has been to discover and amplify understanding of the mechanisms of atherogenic vascular diseases and attendant comorbidities.

GeneSTAR was supported by grants from the National Institutes of Health/National Heart, Lung, and Blood Institute (U01 HL72518, HL087698, HL49762, HL58625, HL071025, HL112064), the

National Institutes of Health/National Institute of Nursing Research (NR0224103), and by a grant from the National Institutes of Health/National Center for Research Resources (M01-RR000052) to the Johns Hopkins General Clinical Research Center. WGS for "NHLBI TOPMed: Genetic Studies of Atherosclerosis Risk" (phs001218) was performed at the Broad Institute of MIT and Harvard (HHSN268201500014C), the Macrogen Corp. (3R01HL112064-04S1), and Illumina (R01HL112064).

Hispanic Community Health Study - Study of Latinos (HCHS/SOL, 2540)

*TOPMed dbGaP accession#: phs001395, Parent dbGaP accession#: phs000810*

The Hispanic Community Health Study / Study of Latinos (HCHS/SOL) is a multi-center epidemiologic study in Hispanic/Latino populations to determine the role of acculturation in the prevalence and development of disease, and to identify risk factors playing a protective or harmful role in Hispanics/Latinos.[36] The goals of the HCHS/SOL include studying the prevalence and development of disease in Hispanics/Latinos, including the role of acculturation, and identifying disease risk factors that play protective or harmful roles in Hispanics/Latinos. A total of 16,415 persons of Cuban, Dominican, Mexican, Puerto Rican, Central American, and South American backgrounds were recruited through four Field Centers affiliated with San Diego State University, Northwestern University in Chicago, Albert Einstein College of Medicine in the Bronx area of New York, and the University of Miami. Seven additional academic centers serve as scientific and logistical support centers. Study participants aged 18-74 years took part in an extensive clinic exam and assessments to ascertain socio-demographic, cultural, environmental and biomedical characteristics. Annual follow-up interviews are conducted to determine a range of health outcomes.

The Hispanic Community Health Study/Study of Latinos was carried out as a collaborative study supported by contracts from the National Heart, Lung, and Blood Institute (NHLBI) to the University of North Carolina (N01-HC65233), University of Miami (N01-HC65234), Albert Einstein College of Medicine (N01-HC65235), Northwestern University (N01-HC65236), and San Diego State University (N01-HC65237). The following Institutes/Centers/Offices contribute to the HCHS/SOL through a transfer of funds to the NHLBI: National Center on Minority Health and Health Disparities, the National Institute of Deafness and Other Communications Disorders, the National Institute of Dental and Craniofacial Research, the National Institute of Diabetes and Digestive and Kidney Diseases, the National Institute of Neurological Disorders and Stroke, and the Office of Dietary Supplements. WGS for "NHLBI TOPMed: Hispanic Community Health Study - Study of Latinos" (phs001395) was performed at the Baylor College of Medicine Human Genome Sequencing Center (HHSN268201600033I).

Hypertension Genetic Epidemiology Network and Genetic Epidemiology Network of Arteriopathy (HyperGEN, 1,797)

*TOPMed dbGaP accession#: phs001293, Parent dbGaP accession#: phs001293*

The Hypertension Genetic Epidemiology Network Study (HyperGEN) - Genetics of Left Ventricular (LV) Hypertrophy is a familial study aimed to understand genetic risk factors for LV hypertrophy by conducting genetic studies of continuous traits from echocardiography exams.[37] The originating HyperGEN study aimed to understand genetic risk factors for hypertension.[38] HyperGEN recruited 470 multiply-affected population-based hypertensive AA sibships (N=1224 siblings) from 1996-1999. HyperGEN probands were ascertained by early onset hypertension (i.e., before 60 years); to participate, they had to have at least one hypertensive sibling who was also willing to participate. Data from detailed clinical exams as well as genotyping data for linkage studies, candidate gene studies and GWAS have been collected and is shared between HyperGEN and the ancillary HyperGEN - Genetics of LV Hypertrophy study.

The HyperGEN Study is part of the National Heart, Lung, and Blood Institute (NHLBI) Family Blood Pressure Program; collection of the data represented here was supported by grants U01 HL054472 (MN Lab), U01 HL054473 (DCC), U01 HL054495 (AL FC), and U01 HL054509 (NC FC). The HyperGEN: Genetics of Left Ventricular Hypertrophy Study was supported by NHLBI grant R01 HL055673 with whole-genome sequencing made possible by supplement -18S1. WGS for "NHLBI TOPMed: Hypertension Genetic Epidemiology Network" (phs001293) was performed at the University of Washington Northwest Genomics Center (3R01HL055673-18S1).

Jackson Heart Study (JHS, 1722)

*TOPMed dbGaP accession#: phs000964, Parent dbGaP accession#: phs000286*

The purpose of the Jackson Heart Study (JHS) is to explore the reasons for heightened cardiovascular disease prevalence among African Americans and to uncover new approaches to reduce it. The JHS is a large, community-based, observational study whose 5,306 participants were recruited from among the non-institutionalized African-American adults from urban and rural areas of the three counties (Hinds, Madison, and Rankin) that make up the Jackson, MS, metropolitan statistical area (MSA).[4,39,40] The JHS design included participants from the Jackson ARIC study who had originally been recruited through random selection from a drivers' license registry. New JHS participants were chosen randomly from the Accudata America commercial listing, which provides householder name, address, zip code, phone number (if available), age group in decades, and family components. In addition, a family component was included in the JHS. The sampling frame for the family study was a participant in any one of the ARIC, random, or volunteer samples whose family size met eligibility requirements. Recruitment was limited to persons 35-84 years old except in the family cohort, where those 21 years old and above were eligible.

The Jackson Heart Study (JHS) is supported and conducted in collaboration with Jackson State University (HHSN268201800013I), Tougaloo College (HHSN268201800014I), the Mississippi State Department of Health (HHSN268201800015I) and the University of Mississippi Medical Center (HHSN268201800010I, HHSN268201800011I and HHSN268201800012I) contracts from the National Heart, Lung, and Blood Institute (NHLBI) and the National Institute on Minority Health and Health Disparities (NIMHD). WGS for "NHLBI TOPMed: The Jackson Heart Study" (phs000964) was performed at the University of Washington

Northwest Genomics Center (HHSN268201100037C). The authors also wish to thank the staffs and participants of the JHS.

Multi-Ethnic Study of Atherosclerosis (MESA, 5,185)

*TOPMed dbGaP accession#: phs001416, Parent dbGaP accession#: phs000209*

The Multi-Ethnic Study of Atherosclerosis (MESA) is a study of the characteristics of subclinical cardiovascular disease (disease detected non-invasively before it has produced clinical signs and symptoms) and the risk factors that predict progression to clinically overt cardiovascular disease or progression of the subclinical disease.[41] MESA researchers study a diverse, population-based sample of 6,814 asymptomatic men and women aged 45-84. Thirty-eight percent of the recruited participants are white, 28 percent African-American, 22 percent Hispanic, and 12 percent Asian, predominantly of Chinese descent. Participants were recruited from six field centers across the United States: Wake Forest University, Columbia University, Johns Hopkins University, University of Minnesota, Northwestern University and University of California - Los Angeles. Each participant received an extensive exam and determination of coronary calcification, ventricular mass and function, flow-mediated endothelial vasodilation, carotid intimal-medial wall thickness and presence of echogenic lucencies in the carotid artery, lower extremity vascular insufficiency, arterial wave forms, electrocardiographic (ECG) measures, standard coronary risk factors, sociodemographic factors, lifestyle factors, and psychosocial factors. Selected repetition of subclinical disease measures and risk factors at follow-up visits allows study of the progression of disease. Blood samples have been assayed for putative biochemical risk factors and stored for case-control studies. DNA has been extracted and lymphocytes cryopreserved (for possible immortalization) for study of candidate genes and possibly, genome-wide scanning, expression, and other genetic techniques. Participants are being followed for identification and characterization of cardiovascular disease events, including acute myocardial infarction and other forms of coronary heart disease (CHD), stroke, and congestive heart failure; for cardiovascular disease interventions; and for mortality.

Whole genome sequencing (WGS) for the Trans-Omics in Precision Medicine (TOPMed) program was supported by the National Heart, Lung and Blood Institute (NHLBI). WGS for “NHLBI TOPMed: Multi-Ethnic Study of Atherosclerosis (MESA)” (phs001416.v1.p1) was performed at the Broad Institute of MIT and Harvard (3U54HG003067-13S1). Centralized read mapping and genotype calling, along with variant quality metrics and filtering were provided by the TOPMed Informatics Research Center (3R01HL-117626-02S1). Phenotype harmonization, data management, sample-identity QC, and general study coordination, were provided by the TOPMed Data Coordinating Center (3R01HL-120393-02S1). MESA and the MESA SHARe project are conducted and supported by the National Heart, Lung, and Blood Institute (NHLBI) in collaboration with MESA investigators. The MESA project is conducted and supported by the National Heart, Lung, and Blood Institute (NHLBI) in collaboration with MESA investigators. Support for MESA is provided by contracts 75N92020D00001, HHSN268201500003I, N01-HC-95159, 75N92020D00005, N01-HC-95160, 75N92020D00002, N01-HC-95161, 75N92020D00003, N01-HC-95162, 75N92020D00006, N01-HC-95163, 75N92020D00004, N01-HC-95164, 75N92020D00007, N01-HC-95165, N01-HC-95166, N01-HC-95167, N01-HC-95168, N01-HC-95169, UL1-TR-000040, UL1-TR-001079, UL1-TR-001420. Support is provided by grants and contracts R01HL071051, R01HL071205, R01HL071250, R01HL071251, R01HL071258, R01HL071259, by the National Center for Research Resources, Grant UL1RR033176. The provision of genotyping data was supported in part by the National Center for Advancing Translational Sciences, CTSI grant UL1TR001881, and the National Institute of Diabetes and Digestive and Kidney Disease Diabetes Research Center (DRC) grant DK063491 to the Southern California Diabetes Endocrinology Research Center.

#### Massachusetts General Hospital Atrial Fibrillation Study (MGH_AF, 682)

*TOPMed dbGaP accession#: phs001062, Parent dbGaP accession#: phs001001*

The Massachusetts General Hospital (MGH) Atrial Fibrillation Study was initiated in 2001.[42,43] The study has enrolled serial probands, unaffected and affected family members with atrial fibrillation. At enrollment participants undergo a structured interview to systematically capture their past medical history, AF treatments, and family history. An electrocardiogram is performed; the results of an echocardiogram are obtained; and blood samples are obtained. For the TOPMed whole genome sequencing project only early-onset atrial fibrillation cases were sequenced. Early-onset atrial fibrillation was defined as an age of onset prior to 66 years of age.

The MGH AF Study was supported by grants to Dr. Ellinor from the Fondation Leducq (14CVD01), the National Institutes of Health to Dr. Ellinor (1RO1HL092577, R01HL128914, K24HL105780) and Dr. Lubitz (1R01HL139731) and by grants from the American Heart Association to Dr. Ellinor (18SFRN34110082) and to Dr. Lubitz (18SFRN34250007). WGS for “NHLBI TOPMed: Massachusetts General Hospital Atrial Fibrillation Study” (phs001062) was performed at the Broad Institute of MIT and Harvard (3R01HL092577-06S1, 3U54HG003067-12S2, 3U54HG003067-13S1, and 3UM1HG008895-01S2)

#### San Antonio Family Study (SAFS, 575)

*TOPMed dbGaP accession#: phs001215, Parent dbGaP accession#: phs000462*

The San Antonio Family Heart Study is a complex pedigree-based mixed longitudinal study designed to identify low frequency or rare variants influencing susceptibility to cardiovascular disease, using whole genome sequence (WGS) information from 3,000 individuals in large Mexican American pedigrees from San Antonio, Texas.[44] The major objectives of this study are to identify low frequency or rare variants in and around known common variant signals for CVD, as well as to find novel low frequency or rare variants influencing susceptibility to CVD. The study began in 1991, and included 1,431 individuals in 42 extended families at baseline. Probands were 40 to 60 year old low-income Mexican Americans selected at random without regard to presence or absence of disease, almost exclusively from Mexican American census tracts in San Antonio, Texas. All first, second, and third -degree relatives of the proband and of the proband's spouse, aged 16 years or above, were eligible to participate in the study. 1,200 WGS at 30X WGS were obtained through Illumina funded by a supplement as part of the NHLBI's TOPMed program.

Collection of the San Antonio Family Study data was supported in part by National Institutes of Health (NIH) grants R01 HL045522, MH078143, MH078111 and MH083824; and whole genome sequencing of SAFS subjects was supported by

U01 DK085524 and R01 HL113323. We are very grateful to the participants of the San Antonio Family Study for their continued involvement in our research programs. WGS for “NHLBI TOPMed: Whole Genome Sequencing to Identify Causal Genetic Variants Influencing CVD Risk - San Antonio Family Studies” (phs001215) was performed at Illumina (3R01HL113323-03S1 and R01HL113322).

Samoan Adiposity Study (Samoan, 1,182)

*TOPMed dbGaP accession#: phs000972, Parent dbGaP accession#: phs000914*

The research goal of the Samoan Adiposity Study is to identify genetic variation that increases susceptibility to obesity and cardiometabolic phenotypes among adult Samoans using genome-wide association (GWAS) methods.[45,46] DNA from peripheral blood and phenotypic information were collected from 3,119 adult Samoans, 23 to 70 years of age. The participants reside throughout the independent nation of Samoa, which is experiencing economic development and the nutrition transition. Genotyping was performed with the Affymetrix Genome-Wide Human SNP 6.0 Array using a panel of approximately 900,000 SNPs. Anthropometric, fasting blood biomarkers and detailed dietary, physical activity, health and socio-demographic variables were collected. Whole genome sequencing of a subset was motivated by the opportunity to create a Samoan-specific reference panel for imputation into the larger parent study.

Data collection was funded by NIH grant R01-HL093093 and R01-HL133040. WGS for “NHLBI TOPMed: Samoan Adiposity Study” (phs000972) was performed at the University of Washington Northwest Genomics Center (HHSN268201100037C and HHSN268201500016C). We thank the Samoan participants of the study and local village authorities. We acknowledge the support of the Samoan Ministry of Health and the Samoa Bureau of Statistics for their support of this research.

Taiwan Study of Hypertension using Rare Variants (THRV, 1,979)

*TOPMed dbGaP accession#: phs001387, Parent dbGaP accession#: phs001387*

The THRV-TOPMed study consists of three cohorts: The SAPPHIRe Family cohort (N=1,271), TSGH (Tri-Service General Hospital, a hospital-based cohort, N=160), and TCVGH (Taichung Veterans General Hospital, another hospital-based cohort, N=922), all based in Taiwan.[47,48] 1,271 subjects were previously recruited as part of the NHLBI-sponsored SAPPHIRe Network (which is part of the Family Blood Pressure Program, FBPP). The SAPPHIRe families were recruited to have two or more hypertensive sibs, some families also with one normotensive/hypotensive sib. The two Hospital-based cohorts (TSGH and TCVGH) both recruited unrelated subjects with different recruitment criteria (matched with SAPPHIRe subjects for age, sex, and BMI category).

The Rare Variants for Hypertension in Taiwan Chinese (THRV) is supported by the National Heart, Lung, and Blood Institute (NHLBI) grant (R01HL111249) and its participation in TOPMed is supported by an NHLBI supplement (R01HL111249-04S1). THRV is a collaborative study between Washington University in St. Louis, LA BioMed at Harbor UCLA, University of Texas in Houston, Taichung Veterans General Hospital, Taipei Veterans General Hospital, Tri-Service General Hospital, National Health Research Institutes, National Taiwan University, and Baylor University. THRV is based (substantially) on the parent SAPPHIRe study, along with additional population-based and hospital-based cohorts. SAPPHIRe was supported by NHLBI grants (U01HL54527, U01HL54498) and Taiwan funds, and the other cohorts were supported by Taiwan funds. WGS for “NHLBI TOPMed: Taiwan Study of Hypertension using Rare Variants” (phs001387) was performed at the Baylor College of Medicine Human Genome Sequencing Center (3R01HL111249-04S1, HHSN26820150015C)

Women’s Health Initiative (WHI, 8,188)

*TOPMed dbGaP accession#: phs001237, Parent dbGaP accession#: phs000200*

The Women's Health Initiative (WHI) is a long-term national health study that has focused on strategies for preventing heart disease, breast and colorectal cancer, and osteoporotic fractures in postmenopausal women (clinicaltrials.gov NCT00000611).[49-51] The original WHI study included 161,808 postmenopausal women enrolled between 1993 and 1998. The Fred Hutchinson Cancer Research Center in Seattle, WA serves as the WHI Clinical Coordinating Center for data collection, management, and analysis of the WHI. The WHI has two major parts: a partial factorial randomized Clinical Trial (CT) and an Observational Study (OS); both were conducted at 40 Clinical Centers nationwide. The CT enrolled 68,132 postmenopausal women between the ages of 50-79 into trials testing three prevention strategies. If eligible, women could choose to enroll in one, two, or all three of the trial components. The components are: hormone therapy trials, dietary modification trial, and calcium / vitamin D trial. The Observational Study (OS) examines the relationship between lifestyle, environmental, medical and molecular risk factors and specific measures of health or disease outcomes. This component involves tracking the medical history and health habits of 93,676 women not participating in the CT. Recruitment for the observational study was completed in 1998 and participants were followed annually for 8 to 12 years.

The WHI program is funded by the National Heart, Lung, and Blood Institute, National Institutes of Health, U.S. Department of Health and Human Services through contracts HHSN268201600018C, HHSN268201600001C, HHSN268201600002C, HHSN268201600003C, and HHSN268201600004C. WGS for “NHLBI TOPMed: Women's Health Initiative” (phs001237) was performed at the Broad Institute of MIT and Harvard (HHSN268201500014C)

UK Biobank (external to TOPMed)

The UK Biobank analyses were conducted using the UK Biobank resource under application 7089.

### AMP-T2D-GENES Consortium

Carlos A. Aguilar-Salinas[1], Gil Atzmon[2,3], Francisco Barajas-Olmos[4], Nir Barzilai[2], John Blangero[5], Michael Boehnke[6], Eric Boerwinkle[7,8], Lori L. Bonnycastle[9], Erwin Bottinger[10,11], Donald W. Bowden[12], Noël P. Burtt[13], Federico Centeno-Cruz[4], John C. Chambers[14-16], Edmund Chan[17], Juliana Chan[18-20], Ching-Yu Cheng[21-23], Yoon Shin Cho[24], Cecilia Contreras-Cubas[4], Emilio J. Córdova[4], Adolfo Correa[25], Ralph A. DeFronzo[26], Ravindranath Duggirala[5], Josée Dupuis[27], Jason A. Flannick[13,28,29], Jose C. Florez[30-33], Ma. Eugenia Garay-Sevilla[34], Humberto García-Ortiz[4], Christian Gieger[35-37], Benjamin Glaser[38], Clicerio Gonzalez[39], Maria Elena Gonzalez-Villalpando[40], Niels Grarup[41], Leif C. Groop[42,43], Myron Gross[44], Christopher Haiman[45], Sohee Han[46], Craig L. Hanis[47], Torben Hansen[41,48], Nancy L. Heard-Costa[49,50], Brian E. Henderson[45], Mi Yeong Hwang[46], Sergio Islas-Andrade[51], Marit E. Jørgensen[52-54], Hyun Min Kang[6], Bong-Jo Kim[46], Young Jin Kim[46], Heikki A. Koistinen[55-57], Jaspal Singh Kooner[58], Johanna Kuusisto[59], Soo Heon Kwak[60], Markku Laakso[59], Leslie A. Lange[61], Jong-Young Lee[62], Juyoung Lee[46], Donna M. Lehman[26], Allan Linneberg[63,64], Jianjun Liu[17,65], Ruth J.F. Loos[66,67], Valeriya Lyssenko[42,68], Ronald C.W. Ma[18-20], Juan Manuel Malacara Hernandez[69], Angélica Martínez-Hernández[4], Mark I. McCarthy[70,71], James B. Meigs[33,72,73], Thomas Meitinger[74,75], Elvia Mendoza-Caamal[4], Karen L. Mohlke[76], Andrew D. Morris[77], Alanna C. Morrison[7], Maggie C.Y. Ng[12,78,79], Peter M. Nilsson[80], Christopher J. O'Donnell[81-84], Lorena Orozco[4], Colin N.A. Palmer[85], Kyong Soo Park[60,86,87], Oluf Pedersen[41], Wendy S. Post[88], Michael Preuss[66], Bruce M. Psaty[89-91], Alexander P. Reiner[92], Cristina Revilla-Monsalve[93], Stephen S. Rich[94], Jerome I. Rotter[95], Danish Saleheen[96,97], Claudia Schurmann[10,11,66], Xueling Sim[98], Rob Sladek[99-101], Kerrin S. Small[102], Wing Yee So[18-20], Xavier Soberón[4], Timothy D. Spector[102], Konstantin Strauch[36,103], Tim M. Strom[75,104], E Shyong Tai[17,98,105], Claudia H.T. Tam[18-20], Yik Ying Teo[98,106,107], Farook Thameem[108], Brian Tomlinson[109], Russell P. Tracy[110,111], Tiinamaija Tuomi[112-114], Jaakko Tuomilehto[115-117], Teresa Tusié-Luna[1,118], Rob M. van Dam[98,119], Ramachandran S. Vasan[49,120], James G. Wilson[121], Daniel R. Witte[122,123], Tien-Yin Wong[21-23]

1 - Instituto Nacional de Ciencias Medicas y Nutricion , Mexico City, Mexico.; 2 - Departments of Medicine and Genetics, Albert Einstein College of Medicine, New York, USA.; 3 - University of Haifa, Faculty of natural science, Haifa, Isarel.; 4 - Instituto Nacional de Medicina Genómica, Mexico City, Mexico.; 5 - Department of Human Genetics and South Texas Diabetes and Obesity Institute, University of Texas Rio Grande Valley School of Medicine, Brownsville, TX 78520, USA.; 6 - Department of Biostatistics and Center for Statistical Genetics, University of Michigan, Ann Arbor, Michigan, USA.; 7 - Human Genetics Center, Department of Epidemiology, Human Genetics, and Environmental Sciences, School of Public Health, The University of Texas Health Science Center at Houston, Houston, TX, USA, 77030.; 8 - Human Genome Sequencing Center, Baylor College of Medicine, Houston, TX, 77030, USA.; 9 - Medical Genomics and Metabolic Genetics Branch, National Human Genome Research Institute, National Institutes of Health, Bethesda, Maryland, USA.; 10 - Hasso Plattner Institute for Digital Health at Mount Sinai, Icahn School of Medicine at Mount Sinai, New York, NY 10029, USA.; 11 - Digital Health Center, Hasso Plattner Institute, University of Potsdam, Potsdam, Germany.; 12 - Department of Biochemistry, Wake Forest School of Medicine, Winston-Salem, NC, USA, 27157.; 13 - Programs in Metabolism and Medical & Population Genetics, Broad Institute, Cambridge, Massachusetts, USA.; 14 - Department of Epidemiology and Biostatistics, Imperial College London, London, UK.; 15 - Imperial College Healthcare NHS Trust, Imperial College London, London, UK.; 16 - Department of Cardiology, Ealing Hospital NHS Trust, Southall, Middlesex, UK.; 17 - Department of Medicine, Yong Loo Lin School of Medicine, National University of Singapore, National University Health System, , Singapore.; 18 - Department of Medicine and Therapeutics, The Chinese University of Hong Kong, Hong Kong, China.; 19 - Hong Kong Institute of Diabetes and Obesity, The Chinese University of Hong Kong, Hong Kong, China.; 20 - Li Ka Shing Institute of Health Sciences, The Chinese University of Hong Kong, Hong Kong, China.; 21 - Department of Ophthalmology, Yong Loo Lin School of Medicine, National University of Singapore, National University Health System, , Singapore.; 22 - Ophthalmology & Visual Sciences Academic Clinical Program (Eye ACP), Duke-NUS Medical School, Singapore.; 23 - Singapore Eye Research Institute, Singapore National Eye Centre, Singapore.; 24 - Department of Biomedical Science, Hallym University, Chuncheon, Republic of Korea.; 25 - Department of Medicine, University of Mississippi Medical Center, Jackson, MS, USA, 39216.; 26 - Department of Medicine, University of Texas Health Science Center, San Antonio, Texas, USA.; 27 - Department of Biostatistics, Boston University School of Public Health, Boston, Massachusetts, USA.; 28 - Department of Pediatrics, Harvard Medical School, Boston, MA, USA, 02115.; 29 - Division of Genetics and Genomics, Boston Children's Hospital, Boston, MA, USA, 02115.; 30 - Center for Genomic Medicine, Department of Medicine, Massachusetts General Hospital, Boston, Massachusetts, USA, 02114.; 31 - Diabetes Research Center (Diabetes Unit), Department of Medicine, Massachusetts General Hospital, Boston, Massachusetts, USA.; 32 - Program in Medical and Population Genetics, Broad Institute of Harvard and MIT, Cambridge, MA, USA, 02142.; 33 - Department of Medicine, Harvard Medical School, Boston, MA, USA, 02115.; 34 - Department of Medical Science, Division of Health Science. University of Guanajuato.; 35 - Research Unit of Molecular Epidemiology, Helmholtz Zentrum München, German Research Center for Environmental Health, Neuherberg, Germany.; 36 - Institute of Epidemiology, Helmholtz Zentrum München, German Research Center for Environmental Health, Neuherberg, Germany.; 37 - German Center for Diabetes Research (DZD), Neuherberg, Germany.; 38 - Endocrinology and Metabolism Service, Hadassah-Hebrew University Medical Center, Jerusalem, Israel.; 39 - Unidad de Diabetes y Riesgo Cardiovascular, Instituto Nacional de Salud Pœblica, Cuernavaca, Morelos, Mexico.; 40 - Centro de Estudios en Diabetes, Mexico City, Mexico.; 41 - Novo Nordisk Foundation Center for Basic Metabolic Research, Faculty of Health and Medical Sciences, University of Copenhagen, Copenhagen, Denmark.; 42 - Department of Clinical Sciences, Diabetes and Endocrinology, Lund University Diabetes Centre, Malmö, Sweden.; 43 - Finnish Institute for Molecular Genetics, University of Helsinki, Helsinki, Finland.; 44 - Department of Laboratory Medicine and Pathology, University of Minnesota, Minneapolis, Minnesota, USA.; 45 - Department of Preventive Medicine, Keck School of Medicine, University of Southern California, Los Angeles, California, USA.; 46 - Division of Genome Science, Department of Precision Medicine, Chungcheongbuk-do, Republic of Korea.; 47 - Human Genetics Center, School of Public Health, The University of Texas Health Science Center at Houston, Houston, Texas, USA.; 48 - Faculty of Health Sciences, University of Southern Denmark, Odense,

Denmark.; 49 - NHLBI Framingham Heart Study, Framingham, Massachusetts, USA.; 50 - Department of Neurology, Boston University School of Medicine, Boston, Massachusetts, USA.; 51 - Dirección de Investigación, Hospital General de México "Dr. Eduardo Liceaga", Secretaría de Salud, Mexico City, Mexico.; 52 - National Institute of Public Health, University of Southern Denmark, Copenhagen, Denmark.; 53 - Steno Diabetes Center Copenhagen, Gentofte, Denmark.; 54 - Greenland Centre for Health Research, University of Greenland, Nuuk, Greenland.; 55 - Department of Public Health Solutions, Finnish Institute for Health and Welfare, Helsinki, Finland.; 56 - Minerva Foundation Institute for Medical Research, Helsinki, Finland.; 57 - University of Helsinki and Department of Medicine, Helsinki University Central Hospital, Helsinki, Finland.; 58 - National Heart and Lung Institute, Cardiovascular Sciences, Hammersmith Campus, Imperial College London, London, UK.; 59 - Institute of Clinical Medicine, University of Eastern Finland, and Kuopio University Hospital, Kuopio, Finland.; 60 - Department of Internal Medicine, Seoul National University Hospital, Seoul, Republic of Korea.; 61 - Department of Medicine, University of Colorado Denver, Anschutz Medical Campus, Aurora, CO, USA 80045.; 62 - Ministry of Health and Welfare, Seoul, Republic of Korea.; 63 - Center for Clinical Research and Prevention, Bispebjerg and Frederiksberg Hospital, The Capital Region, Copenhagen, Denmark.; 64 - Department of Clinical Medicine, Faculty of Health and Medical Sciences, University of Copenhagen, Copenhagen, Denmark.; 65 - Genome Institute of Singapore, Agency for Science, Technology and Research, Singapore.; 66 - Charles R. Bronfman Institute of Personalized Medicine, Icahn School of Medicine at Mount Sinai, New York, New York, USA.; 67 - The Mindich Child Health and Development Institute, Icahn School of Medicine at Mount Sinai, New York, NY 10029.; 68 - University of Bergen, Norway.; 69 - Departments of Medicine and Human Genetics, The University of Chicago, Chicago, Illinois, USA.; 70 - Oxford Centre for Diabetes, Endocrinology and Metabolism, Radcliffe Department of Medicine, University of Oxford, Oxford, UK.; 71 - Wellcome Centre for Human Genetics, University of Oxford, Oxford, UK.; 72 - General Medicine Division, Massachusetts General Hospital, Boston, Massachusetts, USA.; 73 - Broad Institute of MIT and Harvard, Cambridge, Massachusetts, USA.; 74 - Deutsches Forschungszentrum fŸr Herz-Kreislauferkrankungen (DZHK), Partner Site Munich Heart Alliance, Munich, Germany.; 75 - Institute of Human Genetics, Technische Universität München, Munich, Germany.; 76 - Department of Genetics, University of North Carolina, Chapel Hill, North Carolina, USA.; 77 - Clinical Research Centre, Centre for Molecular Medicine, Ninewells Hospital and Medical School, Dundee, UK.; 78 - Center for Diabetes Research, Wake Forest School of Medicine, Winston-Salem, North Carolina, USA.; 79 - Center for Genomics and Personalized Medicine Research, Wake Forest School of Medicine, Winston-Salem, North Carolina, USA.; 80 - Department of Clinical Sciences, Lund University, Malmö, Sweden.; 81 - Section of Cardiology , Department of Medicine, VA Boston Healthcare, Boston, Massachusetts, USA.; 82 - Harvard Medical School, Boston, Massachusetts, USA.; 83 - Department of Medicine, Brigham and Women's Hospital, Boston, MA 02115.; 84 - Intramural Administration Management Branch, National Heart, Lung, and Blood Institute, NIH, Framingham, Massachusetts, USA.; 85 - Pat Macpherson Centre for Pharmacogenetics and Pharmacogenomics,Division of Population Health and Genomics, University of Dundee, Ninewells Hospital and Medical School, Dundee, UK.; 86 - Department of Internal Medicine, Seoul National University College of Medicine, Seoul, Republic of Korea.; 87 - Department of Molecular Medicine and Biopharmaceutical Sciences, Graduate School of Convergence Science and Technology, Seoul National University, Seoul, Republic of Korea.; 88 - Division of Cardiology, Department of Medicine, Johns Hopkins University, Baltimore, Maryland, USA.; 89 - Cardiovascular Health Research Unit, Department of Medicine, University of Washington, Seattle, WA, USA, 98101.; 90 - Department of Epidemiology, University of Washington, Seattle, WA, USA 98101.; 91 - Department of Health Services, University of Washington, Seattle, WA, USA 98101.; 92 - University of Washington, Seattle, Washington, USA.; 93 - Instituto Mexicano del Seguro Social SXXI, Mexico City, Mexico.; 94 - Center for Public Health Genomics, University of Virginia, Charlottesville, VA 22908, USA.; 95 - The Institute for Translational Genomics and Population Sciences, Department of Pediatrics, The Lundquist Institute for Biomedical Innovation at Harbor-UCLA Medical Center, Torrance, CA 90502.; 96 - Department of Biostatistics and Epidemiology, University of Pennsylvania, Philadelphia, Pennsylvania, USA.; 97 - Center for Non-Communicable Diseases, Karachi, Pakistan.; 98 - Saw Swee Hock School of Public Health, National University of Singapore and National University Health System, Singapore.; 99 - Department of Human Genetics, McGill University, Montreal, Quebec, Canada.; 100 - Division of Endocrinology and Metabolism, Department of Medicine, McGill University, Montreal, Quebec, Canada.; 101 - McGill University and Génome Québec Innovation Centre, Montreal, Quebec, Canada.; 102 - Department of Twin Research and Genetic Epidemiology, King's College London, London, UK.; 103 - Institute of Genetic Epidemiology, Helmholtz Zentrum München, German Research Center for Environmental Health, Neuherberg, Germany.; 104 - Institute of Human Genetics, Helmholtz Zentrum München, German Research Center for Environmental Health, Neuherberg, Germany.; 105 - Duke-NUS Medical School Singapore, Singapore.; 106 - Department of Statistics and Applied Probability, National University of Singapore, Singapore.; 107 - Life Sciences Institute, National University of Singapore, Singapore.; 108 - Department of Biochemistry, Faculty of Medicine, Health Science Center, Kuwait University, Safat, Kuwait.; 109 - Faculty of Medicine, Macau University of Science & Technology, Macau, China.; 110 - Department of Biochemistry, Robert Larner, M.D. College of Medicine, University of Vermont, Burlington, Vermont, USA.; 111 - Department of Pathology and Laboratory Medicine, Robert Larner, M.D. College of Medicine, University of Vermont, Burlington, Vermont, USA.; 112 - Department of Endocrinology, Abdominal Centre, Helsinki University Hospital, Helsinki, Finland.; 113 - Folkhälsan Research Centre, Helsinki, Finland.; 114 - Research Programs Unit, Diabetes and Obesity, University of Helsinki, Helsinki, Finland.; 115 - Public Health Promition Unit, Finnish Institute for Health and Welfare, Helsinki, Finland.; 116 - Department of Public Health, University of Helsinki, Helsinki, Finland.; 117 - Diabetes Research Group, King Abdulaziz University, Jeddah, Saudi Arabia.; 118 - Departamento de Medicina Genómica y Toxicología, Instituto de Investigaciones Biomédicas, Universidad Nacional Autónoma de México/ Instituto Nacional de Ciencias Médicas y Nutrición Salvador Zubirán, Mexico City, Mexico.; 119 - Department of Nutrition, Harvard School of Public Health, Boston, Massachusetts, USA.; 120 - Departments of Medicine & Epidemiology, Boston University Schools of Medicine & Public Health, Boston, MA.; 121 - Division of Cardiovascular Medicine, Beth Israel Deaconess Medical Center, Boston, MA 02215.; 122 - Department of Public Health, Aarhus University, Aarhus, Denmark.; 123 - Steno Diabetes Center Aarhus, Aarhus, Denmark.;

### Myocardial Infarction Genetics Consortium

Diego Ardissino[1-3], Usman Baber[4], Matthew J. Bown[5,6], Mark D. Chaffin[7-9], Rajiv Chowdhury[10,11], John Danesh[10,12], Roberto Elosua Llanos[13-15], Valentin Fuster[4,16], Namrata Gupta[7], Sekar Kathiresan[7,8,17,18], Amit V. Khera[7-9], Ruth McPherson[19], Olle Melander[20,21], Marju Orho-Melander[22], Danish Saleheen[23,24], Nilesh Samani[5,6], Heribert Schunkert[25], Minxian X. Wang[7,8,26], Hugh Watkins[27]

1 - ASTC: Associazione per lo Studio Della Trombosi in Cardiologia, Pavia, Italy.; 2 - Azienda Ospedaliero-Universitaria di Parma,ÊParma,Italy.; 3 - Universitˆ degli Studi di Parma, Parma, Italy.; 4 - Department of Medicine, Icahn School of Medicine at Mount Sinai, New York, NY 10029.; 5 - Department of Cardiovascular Sciences, University of Leicester, Leicester, UK.; 6 - NIHR Leicester Biomedical Research Centre, Glenfield Hospital, Leicester UK.; 7 - Program in Medical and Population Genetics, Broad Institute of Harvard and MIT, Cambridge, MA, USA, 02142.; 8 - Center for Genomic Medicine, Department of Medicine, Massachusetts General Hospital, Boston, Massachusetts, USA, 02114.; 9 - Cardiovascular Research Center, Massachusetts General Hospital, Boston, MA, USA, 02114.; 10 - MRC/BHF Cardiovascular Epidemiology Unit, Department of Public Health and Primary Care, University of Cambridge, Cambridge, UK.; 11 - Centre for Non-Communicable disease Research (CNCR), Bangladesh.; 12 - The National Institute for Health Research Blood and Transplant Research Unit (NIHR BTRU) in Donor Health and Genomics at the University of Cambridge, Cambridge, UK.; 13 - Cardiovascular Epidemiology and Genetics, Hospital del Mar Research Institute, Barcelona, Spain.; 14 - CIBER Enfermedades Cardiovasculares (CIBERCV), Barcelona, Spain.; 15 - Facultat de Medicina, Universitat de Vic-Central de Catalu–a, Vic, Spain.; 16 - Centro Nacional de Investigaciones Cardiovasculares Carlos III (CNIC), Madrid, Spain.; 17 - Department of Medicine, Harvard Medical School, Boston, MA, USA, 02115.; 18 - Verve Therapeutics, Cambridge, MA, USA 02139.; 19 - Ruddy Canadian Cardiovascuar Genetics Centre, University of Ottawa Heart Institute, Ottawa, Canada.; 20 - Department of Clinical Sciences, Diabetes and Endocrinology, Lund University Diabetes Centre, Malmö, Sweden.; 21 - Department of Emergency and Internal Medicine, SkŒne University Hospital, Malmö, Sweden.; 22 - Department of Clinical Sciences, Lund University, Malmö, Sweden.; 23 - Department of Biostatistics and Epidemiology, University of Pennsylvania, Philadelphia, Pennsylvania, USA.; 24 - Center for Non-Communicable Diseases, Karachi, Pakistan.; 25 - Deutsches Herzzentrum MŸnchen, Technische UniversitŠt MŸnchen, Deutsches Zentrum fŸr Herz-Kreislauf-Forschung, MŸnchen, Germany.; 26 - Cardiovascular Disease Initaitive, Broad Institute of Harvard and MIT, Cambridge, MA, USA, 02142.; 27 - Cardiovascular Medicine, Radcliffe Department of Medicine and the Wellcome Trust Centre for Human Genetics, University of Oxford, Oxford, UK.;

### NHLBI Trans-Omics for Precision Medicine (TOPMed) Consortium

Namiko Abe[1], Gonalo Abecasis[2], Francois Aguet[3], Christine Albert[4], Laura Almasy[5], Alvaro Alonso[6], Seth Ament[7], Peter Anderson[8], Pramod Anugu[9], Deborah Applebaum-Bowden[10], Kristin Ardlie[3], Dan Arking[11], Donna K Arnett[12], Allison Ashley-Koch[13], Stella Aslibekyan[14], Tim Assimes[15], Paul Auer[16], Dimitrios Avramopoulos[11], John Barnard[17], Kathleen Barnes[18], R. Graham Barr[19], Emily Barron-Casella[11], Lucas Barwick[20], Terri Beaty[11], Gerald Beck[21], Diane Becker[22], Lewis Becker[11], Rebecca Beer[23], Amber Beitelshees[7], Emelia Benjamin[24], Takis Benos[25], Marcos Bezerra[26], Larry Bielak[2], Joshua Bis[27], Thomas Blackwell[2], John Blangero[28], Eric Boerwinkle[29], Donald W. Bowden[30], Russell Bowler[31], Jennifer Brody[8], Ulrich Broeckel[32], Jai Broome[8], Karen Bunting[1], Esteban Burchard[33], Carlos Bustamante[34], Erin Buth[35], Brian Cade[36], Jonathan Cardwell[37], Vincent Carey[38], Cara Carty[39], Richard Casaburi[40], James Casella[11], Peter Castaldi[41], Mark Chaffin[3], Christy Chang[7], Yi-Cheng Chang[42], Daniel Chasman[43], Sameer Chavan[37], Bo-Juen Chen[1], Wei-Min Chen[44], Yii-Der Ida Chen[45], Michael Cho[38], Seung Hoan Choi[3], Lee-Ming Chuang[46], Mina Chung[47], Ren-Hua Chung[48], Clary Clish[49], Suzy Comhair[50], Matthew Conomos[35], Elaine Cornell[51], Adolfo Correa[52], Carolyn Crandall[40], James Crapo[53], L. Adrienne Cupples[54], Joanne Curran[55], Jeffrey Curtis[2], Brian Custer[56], Coleen Damcott[7], Dawood Darbar[57], Sayantan Das[2], Sean David[58], Colleen Davis[8], Michelle Daya[37], Michael DeBaun[59], Dawn DeMeo[38], Ranjan Deka[60], Scott Devine[7], Qing Duan[61], Ravi Duggirala[62], Jon Peter Durda[51], Susan Dutcher[63], Charles Eaton[64], Lynette Ekunwe[9], Adel El Boueiz[65], Patrick Ellinor[66], Leslie Emery[8], Serpil Erzurum[17], Charles Farber[44], Tasha Fingerlin[67], Matthew Flickinger[2], Myriam Fornage[29], Nora Franceschini[68], Chris Frazar[8], Mao Fu[7], Stephanie M. Fullerton[8], Lucinda Fulton[63], Stacey Gabriel[3], Weiniu Gan[23], Shanshan Gao[37], Yan Gao[9], Margery Gass[69], Bruce Gelb[70], Xiaoqi (Priscilla) Geng[2], Mark Geraci[71], Soren Germer[1], Robert Gerszten[72], Auyon Ghosh[38], Richard Gibbs[73], Chris Gignoux[15], Mark Gladwin[25], David Glahn[74], Stephanie Gogarten[8], Da-Wei Gong[7], Harald Goring[75], Sharon Graw[18], Daniel Grine[37], C. Charles Gu[63], Yue Guan[7], Xiuqing Guo[45], Namrata Gupta[3], Jeff Haessler[69], Michael Hall[76], Daniel Harris[7], Nicola L. Hawley[77], Jiang He[78], Ben Heavner[35], Susan Heckbert[8], Ryan Hernandez[33], David Herrington[79], Craig Hersh[80], Bertha Hidalgo[14], James Hixson[29], Brian Hobbs[38], John Hokanson[37], Elliott Hong[7], Karin Hoth[81], Chao (Agnes) Hsiung[82], Yi-Jen Hung[83], Haley Huston[84], Chii Min Hwu[85], Marguerite Ryan Irvin[14], Rebecca Jackson[86], Deepti Jain[8], Cashell Jaquish[23], Min A Jhun[2], Jill Johnsen[84], Andrew Johnson[23], Craig Johnson[8], Rich Johnston[6], Kimberly Jones[11], Hyun Min Kang[87], Robert Kaplan[88], Sharon Kardia[2], Sekar Kathiresan[3], Shannon Kelly[56], Eimear Kenny[70], Michael Kessler[7], Alyna Khan[8], Wonji Kim[89], Greg Kinney[90], Barbara Konkle[84], Charles Kooperberg[69], Holly Kramer[91], Christoph Lange[92], Ethan Lange[37], Leslie Lange[37], Cathy Laurie[8], Cecelia Laurie[8], Meryl LeBoff[38], Jonathon LeFaive[2], Jiwon Lee[38], Seunggeun Shawn Lee[2], Wen-Jane Lee[85], David Levine[8], Dan Levy[23], Joshua Lewis[7], Xiaohui Li[45], Yun Li[61], Henry Lin[45], Honghuang Lin[93], Keng Han Lin[2], Xihong Lin[94], Simin Liu[95], Yongmei Liu[96], Yu Liu[97], Ruth J.F. Loos[98], Steven Lubitz[66], Kathryn Lunetta[93], James Luo[23], Michael Mahaney[55], Barry Make[11], Ani Manichaikul[44], JoAnn Manson[38], Lauren Margolin[3], Lisa Martin[99], Susan Mathai[37], Rasika Mathias[11], Susanne May[35], Patrick McArdle[7], Merry-Lynn McDonald[14], Sean McFarland[89], Stephen McGarvey[64], Daniel McGoldrick[8], Caitlin McHugh[35], Hao Mei[9], Luisa Mestroni[18], Deborah A Meyers[100], Julie Mikulla[23], Nancy Min[9], Mollie Minear[23], Ryan L Minster[25], Braxton D. Mitchell[7], Matt Moll[41], May E. Montasser[7], Courtney Montgomery[101], Arden Moscati[70], Solomon Musani[52], Stanford Mwasongwe[9], Josyf C Mychaleckyj[44], Girish Nadkarni[70], Rakhi Naik[11], Take Naseri[102], Pradeep Natarajan[3], Sergei Nekhai[103], Sarah C. Nelson[35], Bonnie Neltner[37], Deborah Nickerson[8], Kari North[61], Jeff O'Connell[104], Tim O'Connor[7], Heather Ochs-Balcom[105], David T. Paik[106], Nicholette Palmer[107], James Pankow[108], George Papanicolaou[23], Afshin Parsa[7], Juan Manuel Peralta[62], Marco Perez[15], James Perry[7], Ulrike Peters[69], Patricia Peyser[2], Lawrence S Phillips[6], Toni Pollin[7], Wendy Post[109], Julia Powers Becker[110], Meher Preethi Boorgula[37], Michael Preuss[70], Bruce Psaty[8], Pankaj Qasba[23], Dandi Qiao[38], Zhaohui Qin[6], Nicholas Rafaels[111], Laura Raffield[112], Vasan S. Ramachandran [93], D.C. Rao[63], Laura Rasmussen-Torvik[113], Aakrosh Ratan[44], Susan Redline[38], Robert Reed[7], Elizabeth Regan[53], Alex Reiner[114], Muagututia Sefuiva Reupena[115], Ken Rice[8], Stephen Rich[44], Dan Roden[116], Carolina Roselli[3], Jerome Rotter[45], Ingo Ruczinski[11], Pamela Russell[37], Sarah Ruuska[84], Kathleen Ryan[7], Ester Cerdeira Sabino[117], Danish Saleheen[19], Shabnam Salimi[7], Steven Salzberg[11], Kevin Sandow[118], Vijay G. Sankaran[119], Christopher Scheller[2], Ellen Schmidt[2], Karen Schwander[63], David Schwartz[37], Frank Sciurba[25], Christine Seidman[120], Jonathan Seidman[121], Vivien Sheehan[122], Stephanie L. Sherman[123], Amol Shetty[7], Aniket Shetty[37], Wayne Hui-Heng Sheu[85], M. Benjamin Shoemaker[124], Brian Silver[125], Edwin Silverman[38], Jennifer Smith[2], Josh Smith[8], Nicholas Smith[126], Tanja Smith[1], Sylvia Smoller[88], Beverly Snively[127], Michael Snyder[15], Tamar Sofer[38], Nona Sotoodehnia[8], Adrienne M. Stilp[8], Garrett Storm[37], Elizabeth Streeten[7], Jessica Lasky Su[38], Yun Ju Sung[63], Jody Sylvia[38], Adam Szpiro[8], Carole Sztalryd[7], Daniel Taliun[2], Hua Tang[128], Margaret Taub[11], Kent D. Taylor[129], Matthew Taylor[18], Simeon Taylor[7], Marilyn Telen[13], Timothy A. Thornton[8], Machiko Threlkeld[130], Lesley Tinker[69], David

Tirschwell[8], Sarah Tishkoff[131], Hemant Tiwari[132], Catherine Tong[133], Russell Tracy[134], Michael Tsai[108], Dhananjay Vaidya[11], David Van Den Berg[135], Peter VandeHaar[2], Scott Vrieze[108], Tarik Walker[37], Robert Wallace[81], Avram Walts[37], Fei Fei Wang[8], Heming Wang[136], Karol Watson[40], Daniel E. Weeks[25], Bruce Weir[8], Scott Weiss[38], Lu-Chen Weng[66], Jennifer Wessel[137], Cristen Willer[138], Kayleen Williams[35], L. Keoki Williams[139], Carla Wilson[38], James Wilson[140], Quenna Wong[8], Joseph Wu[106], Huichun Xu[7], Lisa Yanek[11], Ivana Yang[37], Rongze Yang[7], Norann Zaghloul[7], Maryam Zekavat[3], Yingze Zhang[141], Snow Xueyan Zhao[53], Wei Zhao[142], Degui Zhi[29], Xiang Zhou[2], Xiaofeng Zhu[143], Michael Zody[1], Sebastian Zoellner[2], Mariza de Andrade[144], Lisa de las Fuentes[145]

1 - New York Genome Center, New York, New York, 10013; 2 - University of Michigan, Ann Arbor, Michigan, 48109; 3 - Broad Institute, Cambridge, Massachusetts, 02142; 4 - Cedars Sinai, Boston, Massachusetts, 02114; 5 - Children's Hospital of Philadelphia, University of Pennsylvania, Philadelphia, Pennsylvania, 19104; 6 - Emory University, Atlanta, Georgia, 30322; 7 - University of Maryland, Baltimore, Maryland, 21201; 8 - University of Washington, Seattle, Washington, 98195; 9 - University of Mississippi, Jackson, Mississippi, 38677; 10 - National Institutes of Health, Bethesda, Maryland, 20892; 11 - Johns Hopkins University, Baltimore, Maryland, 21218; 12 - University of Kentucky, Lexington, Kentucky, 40506; 13 - Duke University, Durham, North Carolina, 27708; 14 - University of Alabama, Birmingham, Alabama, 35487; 15 - Stanford University, Stanford, California, 94305; 16 - University of Wisconsin Milwaukee, Milwaukee, Wisconsin, 53211; 17 - Cleveland Clinic, Cleveland, Ohio, 44195; 18 - University of Colorado Anschutz Medical Campus, Aurora, Colorado, 80045; 19 - Columbia University, New York, New York, 10027; 20 - The Emmes Corporation, LTRC, Rockville, Maryland, 20850; 21 - Cleveland Clinic, Quantitative Health Sciences, Cleveland, Ohio, 44195; 22 - Johns Hopkins University, Medicine, Baltimore, Maryland, 21218; 23 - National Heart, Lung, and Blood Institute, National Institutes of Health, Bethesda, Maryland, 20892; 24 - Boston University, Massachusetts General Hospital, Boston University School of Medicine, Boston, Massachusetts, 02114; 25 - University of Pittsburgh, Pittsburgh, Pennsylvania, 15260; 26 - Fundao de Hematologia e Hemoterapia de Pernambuco - Hemope, Recife, 52011-000; 27 - University of Washington, Cardiovascular Health Research Unit, Department of Medicine, Seattle, Washington, 98195; 28 - University of Texas Rio Grande Valley School of Medicine, Human Genetics, Brownsville, Texas, 78520; 29 - University of Texas Health at Houston, Houston, Texas, 77225; 30 - Wake Forest Baptist Health, Department of Biochemistry, Winston-Salem, North Carolina, 27157; 31 - National Jewish Health, National Jewish Health, Denver, Colorado, 80206; 32 - Medical College of Wisconsin, Milwaukee, Wisconsin, 53226; 33 - University of California, San Francisco, San Francisco, California, 94143; 34 - Stanford University, Biomedical Data Science, Stanford, California, 94305; 35 - University of Washington, Biostatistics, Seattle, Washington, 98195; 36 - Brigham & Women's Hospital, Brigham and Women's Hospital, Boston, Massachusetts, 02115; 37 - University of Colorado at Denver, Denver, Colorado, 80204; 38 - Brigham & Women's Hospital, Boston, Massachusetts, 02115; 39 - Washington State University, Pullman, Washington, 99164; 40 - University of California, Los Angeles, Los Angeles, California, 90095; 41 - Brigham & Women's Hospital, Medicine, Boston, Massachusetts, 02115; 42 - National Taiwan University, Taipei, 10617; 43 - Brigham & Women's Hospital, Division of Preventive Medicine, Boston, Massachusetts, 02215; 44 - University of Virginia, Charlottesville, Virginia, 22903; 45 - Lundquist Institute, Torrance, California, 90502; 46 - National Taiwan University, National Taiwan University Hospital, Taipei, 10617; 47 - Cleveland Clinic, Cleveland Clinic, Cleveland, Ohio, 44195; 48 - National Health Research Institute Taiwan, Miaoli County, 350; 49 - Broad Institute, Metabolomics Platform, Cambridge, Massachusetts, 02142; 50 - Cleveland Clinic, Immunity and Immunology, Cleveland, Ohio, 44195; 51 - University of Vermont, Burlington, Vermont, 05405; 52 - University of Mississippi, Medicine, Jackson, Mississippi, 38677; 53 - National Jewish Health, Denver, Colorado, 80206; 54 - Boston University, Biostatistics, Boston, Massachusetts, 02115; 55 - University of Texas Rio Grande Valley School of Medicine, Brownsville, Texas, 78520; 56 - Vitalant Research Institute, San Francisco, California, 94118; 57 - University of Illinois at Chicago, Chicago, Illinois, 60607; 58 - University of Chicago, Chicago, Illinois, 60637; 59 - Vanderbilt University, Nashville, Tennessee, 37235; 60 - University of Cincinnati, Cincinnati, Ohio, 45220; 61 - University of North Carolina, Chapel Hill, North Carolina, 27599; 62 - University of Texas Rio Grande Valley School of Medicine, Edinburg, Texas, 78539; 63 - Washington University in St Louis, St Louis, Missouri, 63130; 64 - Brown University, Providence, Rhode Island, 02912; 65 - Harvard University, Channing Division of Network Medicine, Cambridge, Massachusetts, 02138; 66 - Massachusetts General Hospital, Boston, Massachusetts, 02114; 67 - National Jewish Health, Center for Genes, Environment and Health, Denver, Colorado, 80206; 68 - University of North Carolina, Epidemiology, Chapel Hill, North Carolina, 27599; 69 - Fred Hutchinson Cancer Research Center, Seattle, Washington, 98109; 70 - Icahn School of Medicine at Mount Sinai, New York, New York, 10029; 71 - Indiana University, Medicine, Indianapolis, Indiana, 46202; 72 - Beth Israel Deaconess Medical Center, Boston, Massachusetts, 02215; 73 - Baylor College of Medicine Human Genome Sequencing Center, Houston, Texas, 77030; 74 - Boston Children's Hospital, Harvard Medical School, Department of Psychiatry, Boston, Massachusetts, 02115; 75 - University of Texas Rio Grande Valley School of Medicine, San Antonio, Texas, 78229; 76 - University of Mississippi, Cardiology, Jackson, Mississippi, 39216; 77 - Yale University, Department of Chronic Disease Epidemiology, New Haven, Connecticut, 06520; 78 - Tulane University, New Orleans, Louisiana, 70118; 79 - Wake Forest Baptist Health, Winston-Salem, North Carolina, 27157; 80 - Brigham & Women's Hospital, Channing Division of Network Medicine, Boston, Massachusetts, 02115; 81 - University of Iowa, Iowa City, Iowa, 52242; 82 - National Health Research Institute Taiwan, Institute of Population Health Sciences, NHRI, Miaoli County, 350; 83 - Tri-Service General Hospital National Defense Medical Center; 84 - Blood Works Northwest, Seattle, Washington, 98104; 85 - Taichung Veterans General Hospital Taiwan, Taichung City, 407; 86 - Ohio State University Wexner Medical Center, Internal Medicine, DIvision of Endocrinology, Diabetes and Metabolism, Columbus, Ohio, 43210; 87 - University of Michigan, Biostatistics, Ann Arbor, Michigan, 48109; 88 - Albert Einstein College of Medicine, New York, New York, 10461; 89 - Harvard University, Cambridge, Massachusetts, 02138; 90 - University of Colorado at Denver, Epidemiology, Aurora, Colorado, 80045; 91 - Loyola University, Public Health Sciences, Maywood, Illinois, 60153; 92 - Harvard School of Public Health, Biostats, Boston, Massachusetts, 02115; 93 - Boston University, Boston, Massachusetts, 02215; 94 - Harvard School of Public Health, Boston, Massachusetts, 02115; 95 - Brown University, Epidemiology and Medicine, Providence, Rhode Island, 02912; 96 - Duke University, Cardiology, Durham, North Carolina, 27708; 97 - Stanford University, Cardiovascular Institute, Stanford, California, 94305; 98 - Icahn School of Medicine at Mount Sinai, The Charles Bronfman Institute for Personalized Medicine, New York, New York, 10029; 99 - George Washington University, Washington, District of Columbia, 20052; 100 - University of Arizona, Tucson, Arizona, 85721; 101 - Oklahoma Medical Research Foundation, Genes and Human Disease, Oklahoma City, Oklahoma, 73104; 102 - Ministry of Health, Government of Samoa, Apia; 103 - Howard University, Washington, District of Columbia, 20059; 104 - University of Maryland, Balitmore, Maryland, 21201; 105 - University at Buffalo, Buffalo, New York, 14260; 106 - Stanford University, Stanford Cardiovascular Institute, Stanford, California, 94305; 107 - Wake Forest Baptist Health, Biochemistry, Winston-Salem, North

Carolina, 27157; 108 - University of Minnesota, Minneapolis, Minnesota, 55455; 109 - Johns Hopkins University, Cardiology/Medicine, Baltimore, Maryland, 21218; 110 - University of Colorado at Denver, Medicine, Denver, Colorado, 80204; 111 - University of Colorado at Denver, Denver, Colorado, 80045; 112 - University of North Carolina, Genetics, Chapel Hill, North Carolina, 27599; 113 - Northwestern University, Chicago, Illinois, 60208; 114 - Fred Hutchinson Cancer Research Center, University of Washington, Seattle, Washington, 98109; 115 - Lutia I Puava Ae Mapu I Fagalele, Apia; 116 - Vanderbilt University, Medicine, Pharmacology, Biomedicla Informatics, Nashville, Tennessee, 37235; 117 - Universidade de Sao Paulo, Faculdade de Medicina, Sao Paulo, 01310000; 118 - Lundquist Institute, TGPS, Torrance, California, 90502; 119 - Harvard University, Division of Hematology/Oncology, Cambridge, Massachusetts, 02138; 120 - Harvard Medical School, Genetics, Boston, Massachusetts, 02115; 121 - Harvard Medical School, Boston, Massachusetts, 02115; 122 - Baylor College of Medicine, Pediatrics, Atlanta, Georgia, 30307; 123 - Emory University, Human Genetics, Atlanta, Georgia, 30322; 124 - Vanderbilt University, Medicine/Cardiology, Nashville, Tennessee, 37235; 125 - UMass Memorial Medical Center, Worcester, Massachusetts, 01655; 126 - University of Washington, Epidemiology, Seattle, Washington, 98195; 127 - Wake Forest Baptist Health, Biostatistical Sciences, Winston-Salem, North Carolina, 27157; 128 - Stanford University, Genetics, Stanford, California, 94305; 129 - Lundquist Institute, Institute for Translational Genomics and Populations Sciences, Torrance, California, 90502; 130 - University of Washington, University of Washington, Department of Genome Sciences, Seattle, Washington, 98195; 131 - University of Pennsylvania, Genetics, Philadelphia, Pennsylvania, 19104; 132 - University of Alabama, Biostatistics, Birmingham, Alabama, 35487; 133 - University of Washington, Department of Biostatistics, Seattle, Washington, 98195; 134 - University of Vermont, Pathology & Laboratory Medicine, Burlington, Vermont, 05405; 135 - University of Southern California, USC Methylation Characterization Center, University of Southern California, California, 90033; 136 - Brigham & Women's Hospital, Mass General Brigham, Boston, Massachusetts, 02115; 137 - Indiana University, Epidemiology, Indianapolis, Indiana, 46202; 138 - University of Michigan, Internal Medicine, Ann Arbor, Michigan, 48109; 139 - Henry Ford Health System, Detroit, Michigan, 48202; 140 - Beth Israel Deaconess Medical Center, Cardiology, Cambridge, Massachusetts, 02139; 141 - University of Pittsburgh, Medicine, Pittsburgh, Pennsylvania, 15260; 142 - University of Michigan, Department of Epidemiology, Ann Arbor, Michigan, 48109; 143 - Case Western Reserve University, Department of Population and Quantitative Health Sciences, Cleveland, Ohio, 44106; 144 - Mayo Clinic, Health Sciences Research, Rochester, Minnesota, 55905; 145 - Washington University in St Louis, Department of Medicine, Cardiovascular Division, St. Louis, Missouri, 63110
